# Reconstitution of chromatin reorganization during mammalian oocyte development

**DOI:** 10.1101/2024.07.27.605426

**Authors:** Jing Wang, Wang Li, Jing Guo, Xiaoming Xu, Ge Lin, Bin Li, Chun So

## Abstract

**Significance:** The generation of developmentally competent eggs requires oocyte growth and maturation. When compared to meiotic maturation, relatively little is known about the key processes that happen in growing oocytes. Since the first visualization of oocyte chromatin under the electron microscope, we have known that growing oocytes undergo a fundamental transition from the non-surrounded nucleolus (NSN) to surrounded nucleolus (SN) chromatin configuration. NSN-to-SN transition is crucial for subsequent embryonic development after fertilization, but what drives NSN-to-SN transition during mammalian oocyte development is still unknown. Here we show that natural degradation of RNA polymerase II (RNAPII) by segregase and proteasome drives NSN-to-SN transition, challenging the conventional view that transcriptional silencing and NSN-to-SN transition are driven by independent epigenetic mechanisms in mammalian oocytes.

The final step of oocyte growth, which reorganizes chromatin from the uncondensed, non-surrounded nucleolus (NSN) configuration to the condensed, surrounded nucleolus (SN) configuration, is essential for embryonic development after fertilization. However, the mechanism of NSN-to-SN transition remains unknown. Here, we identify RNA polymerase II (RNAPII) degradation as the key driver of this process. RNAPII inhibitors, but not nucleoside-based transcription inhibitors, swiftly induced RNAPII degradation and NSN-to-SN transition in oocytes. Strikingly, induced SN-like nuclei recapitulated epigenetic features, chromatin interactions and developmental potential in SN oocytes. Using segregase and proteasome inhibitors and the newly established scFv miniTrim-Away for nuclear proteins, we further demonstrate that RNAPII degradation is necessary and sufficient for NSN-to-SN transition in mouse and human. RNAPII degradation globally cleared RNAPII from transcriptionally active chromatin and increased chromatin dynamics, concomitantly leading to transcriptional silencing and chromatin rearrangement in oocytes. Our study elucidates the framework of chromatin reorganization in growing mammalian oocytes and provides an alternative source of fully grown oocyte nuclei, representing a milestone for the potential use of growing oocytes in reproductive medicine.

---

Oogenesis is essential for sexual reproduction in mammals. Growing oocytes have a non-surrounded nucleolus (NSN) chromatin configuration and actively transcribe mRNAs^1,2^, which are used to establish protein and mRNA stores for the next generation. During the final stages of oogenesis, condensed chromatin forms a ring around the nucleolus in a surrounded nucleolus (SN) configuration and transcription ceases^1,2^. SN oocytes have a higher developmental competence than NSN oocytes^3–8^, and their nucleus primarily accounts for their differences in the success of embryonic development^9^. Therefore, the reorganization of chromatin from NSN to SN configuration is tied to the developmental competence of mammalian oocytes.

NSN-to-SN transition is accompanied by many characteristic changes in the nuclear architecture^10^. Compared to NSN nuclei, SN nuclei display a reduction in nucleolar components and rRNA synthesis, clustering of centromeres and pericentromeric constitutive heterochromatin (chromocenters) around the nucleolus, fewer but larger splicing-related nuclear speckles and altered histone modifications and DNA methylations.

How the chromatin is reorganized during this key step of mammalian oocyte development is a long-standing question. Previous studies of knockout mice have reported a number of proteins that could alter the NSN/SN ratio of isolated oocytes^11–21^, but whether these proteins directly drive NSN-to-SN transition is not known. Furthermore, the physiological relevance of these proteins is unclear: evidence that their expression or activity changes during NSN-to-SN transition in vivo is lacking. Here, we comprehensively mapped almost 30 nuclear components in NSN and SN oocytes to uncover the underlying nuclear changes and to identify candidates that drive the physiological NSN-to-SN transition.

## Results

### Comprehensive analysis of NSN and SN nuclei

First, we established a workflow for studying NSN and SN oocytes by sorting freshly isolated germinal vesicle (GV) oocytes based on their chromatin configurations, followed by fixation, immunofluorescence staining, high-resolution microscopy, and machine learning-based 3D segmentation analyses (Figure 1A). To avoid phototoxicity and DNA damage incurred by the use of UV-excitable Hoechst dyes, we performed live staining of oocyte chromatin using the bright green-excitable SPY555-DNA or red-excitable 5-SiR-Hoechst dyes, which are routinely used for long-term live imaging of human oocytes^22^. NSN and SN oocytes were sorted with a high accuracy of 99.0% and 100%, respectively (Figure 1B). 3D segmentation analysis confirmed that the chromatin volume of NSN and SN oocytes differed by two-fold (Figure 1C), consistent with chromatin condensation^23^ and detachment from the nuclear envelope^24^ during NSN-to-SN transition.

**Figure 1.**
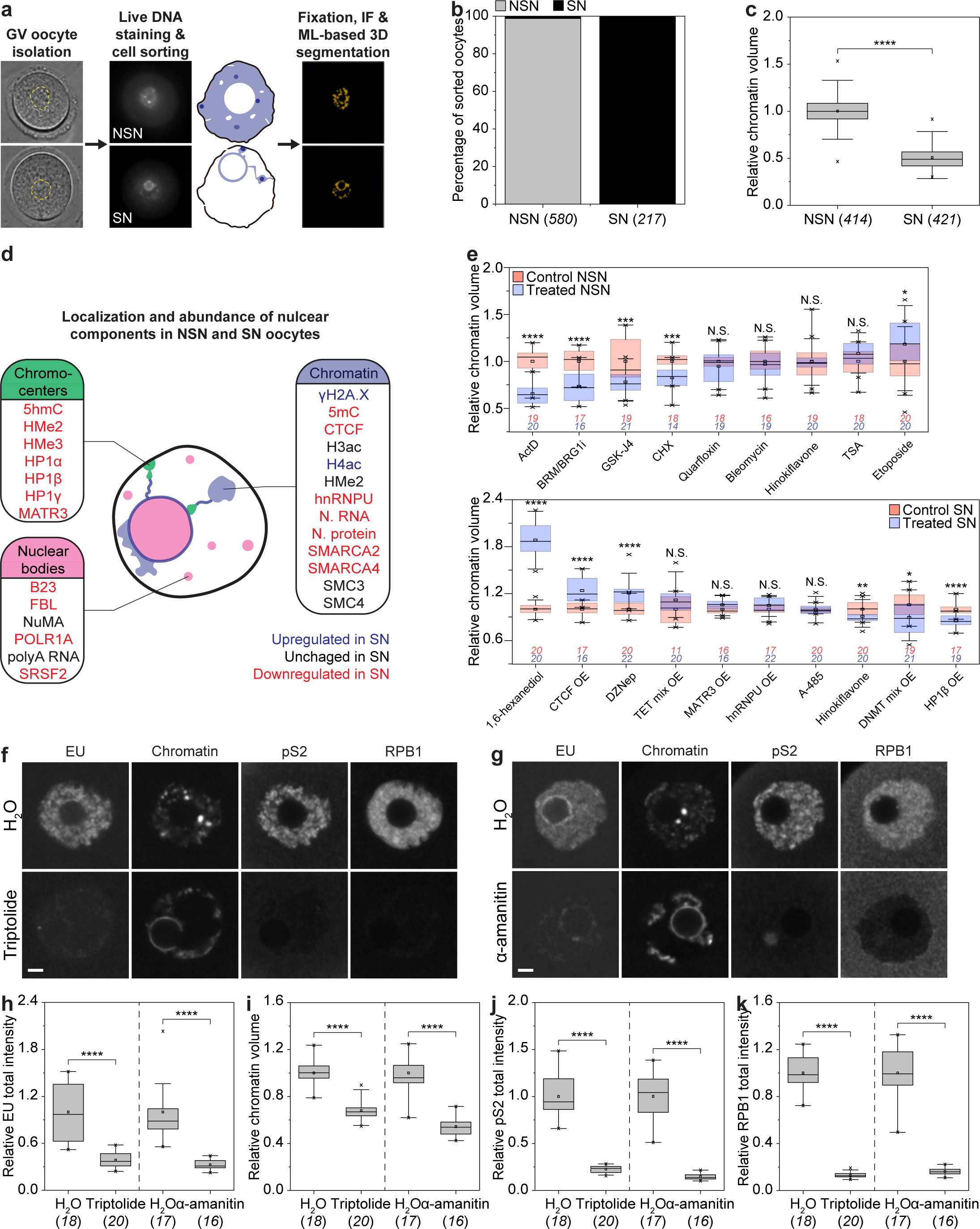
RNAPII inhibitors ectopically induce RNAPII degradation and NSN-to-SN transition in oocytes. **(A)** Workflow for live sorting of NSN and SN oocytes and 3D segmentation analyses of oocyte nuclei. **(B)** Manual scoring of chromatin configurations in sorted NSN and SN oocytes after fixation and staining with Hoechst. **(C)** Automated quantification of chromatin volume in NSN and SN oocytes. **(D)** Schematic representation of oocyte nucleus (scheme: chromocenters in green, nuclear bodies in magenta, and chromatin in blue). **(E)** Automated quantification of chromatin volume in NSN and SN oocytes treated with different small molecule-inhibitors or overexpressing different nuclear components. **(F and G)** Immunofluorescence images of NSN oocytes treated with triptolide or α-amanitin. **(H to K)** Automated quantification of total intensity of EU, chromatin volume, total intensity of pS2 and RPB1 in NSN oocytes treated with triptolide or α-amanitin. *P < 0.05. **P < 0.01. ***P < 0.001. ****P < 0.0001. N.S., not significant. See Methods for box plot specifications. The number of oocytes analyzed is specified in italics. Scale bars are 5 µm.

To study the molecular differences between NSN and SN nuclei, we systematically evaluated the localization and abundance of 28 common nuclear components (Figure 1D, S1A-S1D). We found that 13 of these nuclear components localized to chromatin, and most of them were downregulated in SN nuclei. Seven nuclear components localized to chromocenters and were all downregulated in SN nuclei. Nucleolar proteins and the nuclear speckle marker SRSF2 were downregulated in SN nuclei, in line with previous reports^25^. Thus, most nuclear components are downregulated during NSN-to-SN transition.

### RNA polymerase II (RNAPII) inhibitors ectopically induce NSN-to-SN transition in oocytes

NSN-to-SN transition did not spontaneously occur in vitro even after prolonged culture of NSN oocytes (Figures S2A-S2C), in line with early studies^3,26^. In order to identify nuclear components potentially driving NSN-to-SN transition, we used small-molecule inhibition and overexpression approaches to transiently modulate the levels or activities of different nuclear components, thereby mimicking their differences in NSN and SN oocytes (Figures 1D and S1A). Of almost 65 different treatments (Table S1), only 18 of them led to the desired changes of nuclear components (Figure S2D). Many of these treatments altered the transcriptional activity and/or chromatin compaction in NSN and SN oocytes (Figures 1E and S2E). In particular, treating NSN oocytes with ActD, BRM/BRG1i or GSK-J4 simultaneously silenced transcription and compacted chromatin, whereas CTCF overexpression or DZNep treatment of SN oocytes simultaneously activated transcription and decompacted chromatin. Although no treatment was able to fully reverse NSN-to-SN transition, ActD was able to induce NSN-to-SN transition as evidenced by the ring-like chromatin configuration around the nucleolus and the almost two-fold reduction in chromatin volume (Figures 1E, S2F and S2G).

We also considered the contribution of liquid-liquid phase separation (LLPS), which has recently been implicated in driving chromatin reorganization in sperm cells of flowering plants^27^. To test if LLPS drives NSN-to-SN transition, we treated SN oocytes with 1,6-hexanediol, which disrupts LLPS by weakening hydrophobic interactions between proteins. Although 1,6-hexanediol drastically decompacted chromatin in SN oocytes (Figure 1E), the morphology of chromatin did not resemble that of NSN oocytes, as characterized by the excessive fragmentation of chromocenters (Figure S2G). Also, 1,6-hexanediol collapsed the spherical nucleolus in oocytes (Figure S2G). Despite the absence of the canonical ring-like configuration, chromatin remained tightly associated with the irregularly shaped remnant of the nucleolus (Figure S2G). These observations suggest that the nucleolus provided a rigid surface for the attachment of condensed chromatin during NSN-to-SN transition, in line with the absence of SN configuration but the presence of condensed and transcriptionally silent chromatin in oocytes from nucleolus-deficient mice^28,29^.

ActD acts on both RNAPI and RNAPII^30^. The RNAPI inhibitor quarfloxin did not phenocopy ActD (Figure 1E), suggesting that NSN-to-SN transition involves RNAPII. To test this, we treated NSN oocytes with two widely used RNAPII inhibitors, triptolide and α-amanitin, which covalently binds to the general transcription factor TFIIH to block transcription initiation and impairs translocation to block transcription elongation^30^, respectively. As expected, both triptolide and α-amanitin almost completely abolished nascent transcription in NSN oocytes (Figures 1F-1H). Importantly, both treatments induced NSN-to-SN transition (Figures 1F, 1G and 1I), as supported by live-cell imaging (Figure S3; Videos S1-S4). Thus, RNAPII inhibitors swiftly induce NSN-to-SN transition in oocytes.

### Induced SN-like nuclei recapitulate epigenetic features, chromatin interactions and developmental potential in SN oocytes

To confirm that RNAPII inhibitors induce NSN-to-SN transition beyond morphological changes, we also examined changes in epigenetic features, chromatin interactions and developmental potential in induced SN-like nuclei. DNA damage marker γH2A.X and pan-histone H4 acetylation (H4ac) were upregulated, whereas DNA methylation, heterochromatin protein HP1β, DNA hydroxymethylation and pan-histone dimethylation (HMe2) were downregulated in α-amanitin-treated NSN oocytes (Figures 2A and 2B), similar to SN oocytes (Figures 1D and S1A-S1D). Low-input Hi-C profiling^31^ further revealed that the chromatin contact map from α-amanitin-treated NSN oocytes is more similar to SN oocytes as compared to NSN oocytes (Figure 2C). Moreover, α-amanitin-treated NSN oocytes and SN oocytes had almost identical contact probability curves with significantly more long-range (>1 Mb) contacts than NSN oocytes (Figure 2D), in line with previous reports^23^. Strikingly, in vitro fertilization (IVF) after in vitro maturation (IVM) of α-amanitin-treated NSN oocytes yielded twice more 2-cell embryos as compared to NSN oocytes (Figure 2E). Thus, induced SN-like nuclei recapitulate epigenetic features, chromatin interactions and developmental potential in SN oocytes.

**Figure 2.**
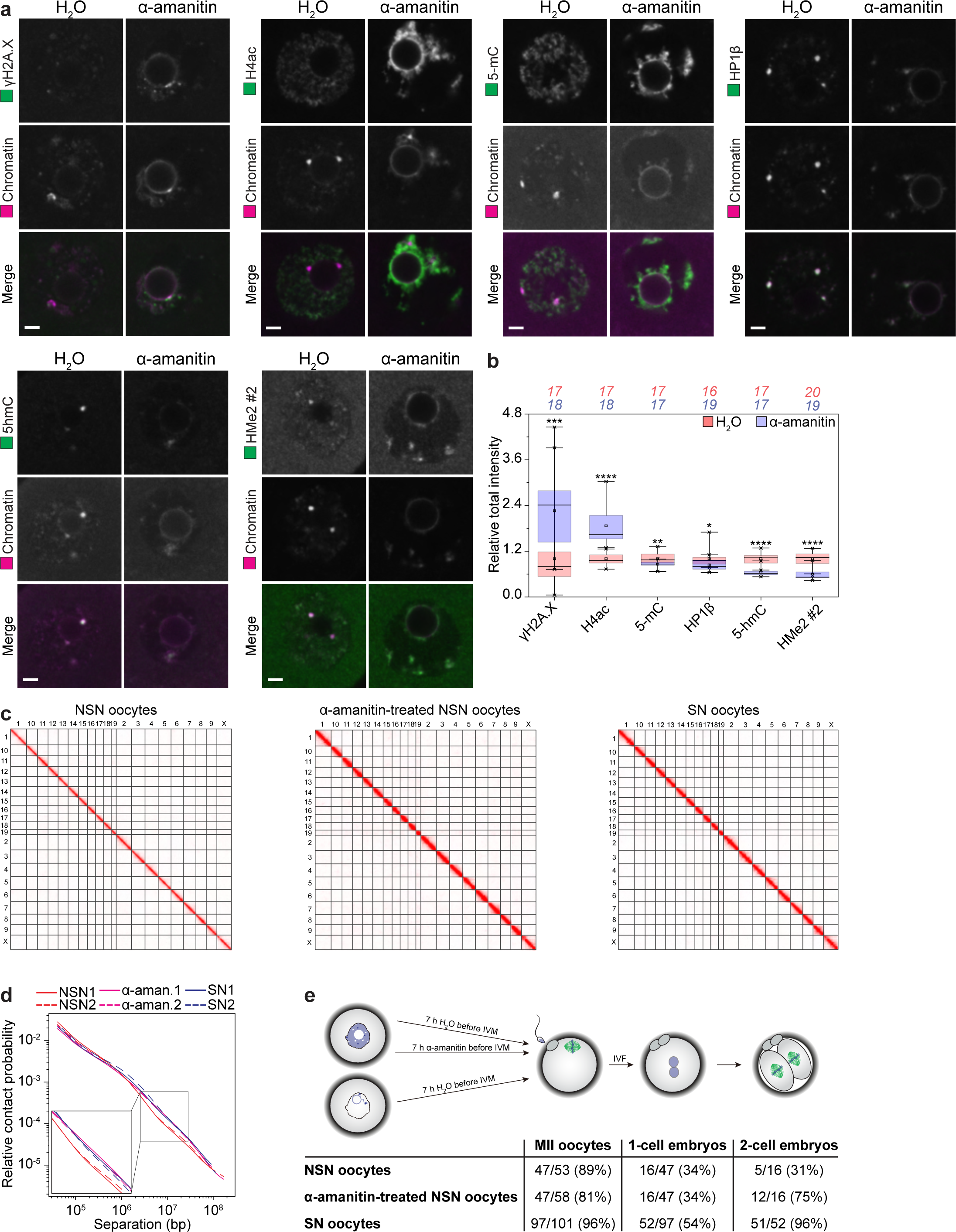
Induced SN-like nuclei recapitulate epigenetic features, chromatin interactions and developmental potential in SN oocytes. **(A)** Immunofluorescence images of NSN oocytes treated with α-amanitin. Green, protein of interest; magenta, chromatin (Hoechst). **(B)** Automated quantification of total intensity of different nuclear components in NSN oocytes treated with α-amanitin. **(C)** Chromatin contact maps from NSN, α-amanitin-treated NSN and SN oocytes. **(D)** Contact probability curves in NSN, α-amanitin-treated NSN and SN oocytes. **(E)** Manual scoring of IVM and IVF of NSN, α-amanitin-treated NSN and SN oocytes. (Fisher’s exact test for 2-cell embryos from NSN and α-amanitin-treated NSN oocytes, P = 0.032). *P < 0.05. **P < 0.01. ***P < 0.001. ****P < 0.0001. See Materials and Methods for box plot specifications. The number of oocytes analyzed is specified in italics. Scale bars are 5 µm.

### Transcriptional silencing is insufficient to induce NSN-to-SN transition in oocytes

Apart from direct inhibition, studies in other systems reported that triptolide and α-amanitin can also trigger RNAPII degradation^32,33^. We performed immunofluorescence staining with a monoclonal antibody against phospho-serine 2 (pS2) at the C-terminal repeat domain and with a monoclonal antibody against the N-terminal domain of the largest RNAPII subunit RPB1, and found that both phosphorylated and total RNAPII were indeed degraded in triptolide- and α-amanitin-treated NSN oocytes (Figures 1F, 1G, 1J and 1K).

To determine if transcriptional silencing alone is sufficient to induce NSN-to-SN transition, we prematurely terminated transcription by incorporating modified adenosine analogues such as 8-aminoadenosine (8-AA) and tubercidin/7-deazaadenosine (7-DAA)^34^. Both 8-AA and 7-DAA suppressed nascent transcription to a level that was similar to triptolide and α-amanitin treatments (Figures S4A-S4C), but without degrading RNAPII (Figures S4A, S4B, S4D and S4E). Importantly, however, 8-AA and 7-DAA did not induce NSN-to-SN transition (Figures S4A, S4B and S4F). Thus, transcriptional silencing is not sufficient to induce NSN-to-SN transition in oocytes.

### Targeted degradation of RNAPII is sufficient to induce NSN-to-SN transition in mouse and human oocytes

Our data suggest that RNAPII degradation might be critical for the NSN-to-SN transition induced by triptolide and α-amanitin. Because RNAPII is an essential gene and oocyte-specific Cre lines all express Cre during early stages of oogenesis, precluding conditional knockout analysis, we aimed to test this hypothesis by acutely degrading RNAPII in NSN oocytes using Trim-Away, an endogenous protein degradation technique based on TRIM21-mediated proteasomal degradation of antibody-antigen complexes^35^. Trim-Away has been used for targeted degradation of cytoplasmic proteins^35^, but RNAPII is a nuclear protein that does not shuttle in and out of the nucleus, so we sought to use the single-chain fragment variable (scFv) for Trim-Away in the nucleus. To this end, we initially tested a scFv developed against pS2 (clone 42B3)^36^ by expressing a GFP-fusion protein in oocytes. Although the 42B3-GFP construct did not localize to the nucleus (Figure S5A), a 42B3-split GFP construct was able to enter the nucleus and colocalize with pS2 on the chromatin in oocytes (Figure S5B). These data show that the size of fusion tag could interfere with the nuclear localization of scFv. Hence, we fused 42B3 with the RING domain of Trim21 (t21R)^37^ rather than the Fc domain of IgG^35^ to develop an scFv miniTrim-Away approach for targeted degradation of nuclear proteins (Figure 3A).

**Figure 3.**
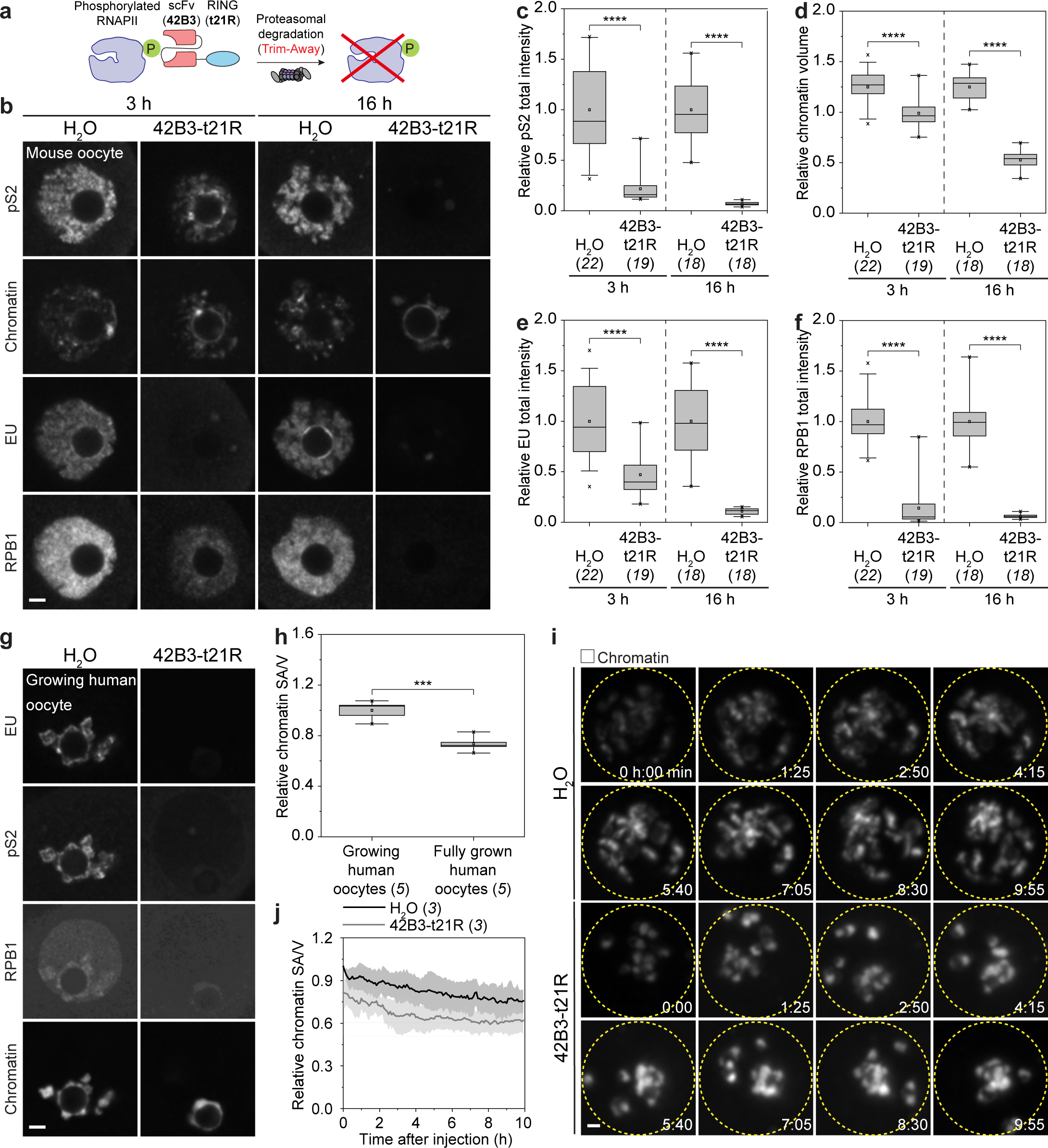
Targeted degradation of RNAPII is sufficient to induce NSN-to-SN transition in mouse and human oocytes. **(A)** Schematic representation of the scFv miniTrim-Away approach. **(B)** Immunofluorescence images of NSN oocytes expressing 42B3-t21R to deplete phosphorylated RNAPII for 3 or 16 h. **(C to F)** Automated quantification of total intensity of pS2, chromatin volume, total intensity of EU and RPB1 in NSN oocytes expressing 42B3-t21R to deplete phosphorylated RNAPII for 3 or 16 h. **(G)** Immunofluorescence images of growing human oocytes expressing 42B3-t21R to deplete phosphorylated RNAPII for 16 h. **(H)** Automated quantification of chromatin surface area-to-volume ratio (SA/V) in growing and fully grown human oocytes. **(I)** Still images from time-lapse movies of growing human oocytes expressing 42B3-t21R. Time is given as hours:minutes from the start of imaging immediately after injection. Yellow dotted circles highlight the nuclei. **(J)** Automated quantification of chromatin SA/V in growing human oocytes expressing 42B3-t21R over time. ***P < 0.001. ****P < 0.0001. N.S., not significant. See Materials and methods for box plot specifications. The number of oocytes analyzed is specified in italics. Scale bars are 5 µm.

Expression of 42B3-t21R in NSN oocytes depleted not only pS2 (Figures 3B and 3C), but also RNAPII phosphorylated at other residues at the C-terminal repeat domain (Figure S5C) and unphosphorylated RNAPII (Figure S5D). These data suggest that RNAPII dynamically cycles between unphosphorylated and different phosphorylated states, consistent with active transcription in NSN oocytes. NSN oocytes expressing 42B3-t21R for 3 hours underwent a partial NSN-to-SN transition, as supported by the 1.5-fold reduction in chromatin volume (Figures 3B and 3D). The presence of residual nascent transcription (Figures 3B and 3E) hinted that the partial NSN-to-SN transition could be due to the residual RNAPII (Figures 3B and 3F), so we extended the expression time from 3 to 16 hours. Remarkably, NSN oocytes expressing 42B3-t21R for 16 hours underwent a full NSN-to-SN transition, as confirmed by the 2-fold reduction in chromatin volume and live-cell imaging (Figures 3B, 3D, S5E and S5F; Videos S5 and S6).

We also tested whether RNAPII degradation drives NSN-to-SN transition in human oocytes. Because human oocytes with an NSN configuration were rarely found (1/52) in immature GV oocytes leftover from fertility treatment, we utilized the growing ones that had partially degraded phosphorylated RNAPII and partially compacted chromatin (Figure 3G) and specifically used surface area-to-volume (SA/V) ratio instead of volume to quantify the less drastic changes in chromatin compaction (Figure 3H). 42B3-t21R was able to fully deplete RNAPII in growing human oocytes (Figure 3G), and we found that this treatment triggered NSN-to-SN transition (Figure 3I; Videos S7 and S8), as supported by the reduction in chromatin SA/V ratio (Figure 3J). Thus, targeted degradation of RNAPII is sufficient to induce NSN-to-SN transition in mouse and human oocytes.

### Segregase- and proteasome-mediated degradation of RNAPII is necessary for NSN-to-SN transition in vivo

Next, we asked whether RNAPII degradation is physiologically relevant to NSN-to-SN transition in vivo. Previous work reported that the levels of phosphorylated RNAPII are lower in SN oocytes as compared to NSN oocytes^38–47^, but whether this observation reflects simply dephosphorylation or depletion at protein level is not clear. To this end, we compared the abundance of phosphorylated as well as of total RNAPII in NSN and SN oocytes by immunofluorescence staining and immunoblotting. We found that phosphorylated RNAPII is not dephosphorylated but specifically depleted (Figures S6A-S6D), accounting for the substantial reduction in total RNAPII levels in SN oocytes (Figures S6E-S6G). As an orthogonal approach, we cultured mouse ovarian follicles ex vivo and harvested oocytes at different growth stages. This approach showed that phosphorylated RNAPII was loaded during the early growth stages, but then progressively depleted during the late growth stages (Figures 4A-4D), concomitant with NSN-to-SN transition (Figures 4A and 4E). The levels of phosphorylated RNAPII in oocytes were not restored until meiotic resumption and fertilization (Figure S6H). We also examined human follicles from ovarian cortical tissues and found similar changes in phosphorylated RNAPII levels (Figures 4F-4H), suggesting that phosphorylated RNAPII was depleted during NSN-to-SN transition in mouse and human.

**Figure 4.**
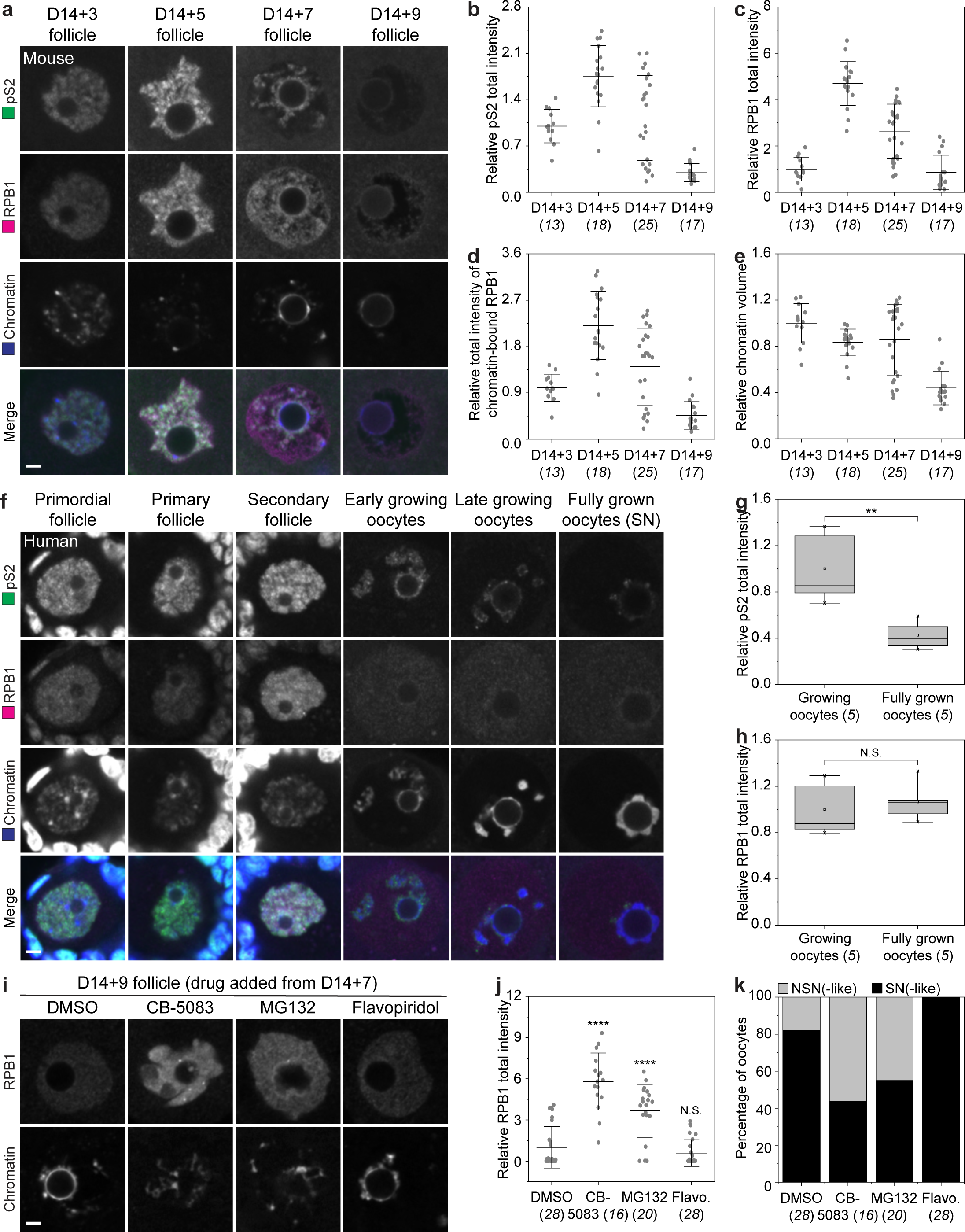
RNAPII degradation is necessary for NSN-to-SN transition in vivo. **(A)** Immunofluorescence images of growing oocytes harvested from mouse secondary follicles cultured for different days. Green, pS2; magenta, RPB1; blue, chromatin (Hoechst). **(B to E)** Automated quantification of total intensity of pS2, RPB1, chromatin-bound RPB1 and chromatin volume in growing oocytes harvested from mouse secondary follicles cultured for different days. **(F)** Immunofluorescence images of human follicles and oocytes of different growth stages. Green, pS2; magenta, RPB1; blue, chromatin (Hoechst). **(G and H)** Automated quantification of total intensity of pS2 and RPB1 in growing and fully grown human oocytes. **(I)** Immunofluorescence images of growing oocytes treated with CB-5083, MG132 and flavopiridol. **(J)** Automated quantification of total intensity of RPB1 in growing oocytes treated with CB-5083, MG132 or flavopiridol. **(K)** Manual scoring of chromatin configurations in growing oocytes treated with CB-5083 (Fisher’s exact test, P = 0.0169), MG132 (Fisher’s exact test, P = 0.0569) or flavopiridol. **P < 0.01. ****P < 0.0001. N.S., not significant. See Materials and methods for box plot specifications. The number of oocytes analyzed is specified in italics. Scale bars are 5 µm.

To decipher how RNAPII levels decrease during NSN-to-SN transition, we first examined the transcriptional regulation of *Rpb1*. Briefly, we performed low-input whole-genome bisulfite sequencing (WGBS)^48^, ATAC-seq^49^ and fractionated RNA-seq and found that DNA methylation levels, chromatin accessibility, and actively transcribing and stored mRNA levels of *Rbp1* were not different between NSN and SN oocytes (Figure S7A). Consistent with our multiomics analyses, RT-qPCR confirmed that *Rpb1* mRNA levels were only marginally reduced in SN oocytes as compared to NSN oocytes (Figure S7B). We then examined the translational regulation of *Rpb1* by Ribo-seq^50^, poly(A) tail-length assay, RIBOmap^51^ and CHX treatment. *Rpb1* mRNA did not display a significant difference in ribosome occupancy nor poly(A) tail-length (Figures S7A and S7C). In addition, RIBOmap analysis and CHX treatment confirmed that *Rpb1* mRNA was not actively translated in NSN or SN oocytes (Figures S7D-S7F). These data suggest that the synthesis of RNAPII protein does not change during NSN-to-SN transition.

To determine whether protein degradation leads to the depletion of phosphorylated RNAPII during NSN-to-SN transition in vivo, we treated cultured mouse ovarian follicles with inhibitors against segregase and proteasome, two key players of proteolysis in cells. Both segregase inhibitor CB-5083 and proteasome inhibitor MG132 suppressed RNAPII degradation (Figures 4I and 4J). Strikingly, they also blocked NSN-to-SN transition (Figures 4I and 4K). Interestingly, however, inhibiting RNAPII phosphorylation with the pan-CDK inhibitor flavopiridol had no effects on RNAPII degradation (Figures 4I and 4J), suggesting that phosphorylation marks the fraction of RNAPII to be degraded but does not mediate the degradation process. Taken together, RNAPII degradation is necessary for NSN-to-SN transition in vivo.

### RNAPII degradation globally clears RNAPII from transcriptionally active chromatin and increases chromatin dynamics during NSN-to-SN transition

To understand further how RNAPII degradation induces NSN-to-SN transition, we investigated the chromatin binding patterns of RNAPII in oocytes. High-resolution imaging revealed that phosphorylated RNAPII is found on chromatin in NSN oocytes, whereas unphosphorylated RNAPII is found in the nucleoplasm in both NSN and SN oocytes (Figure 5A), suggesting that RNAPII degradation removes primarily chromatin-bound RNAPII during NSN-to-SN transition. By optimizing low-input CUT&Tag^52^ (Figure S8A) to map the chromatin binding profile of phosphorylated RNAPII in NSN oocytes, we found that phosphorylated RNAPII was bound to different chromosomes (Figure S8B) and enriched at the promoter, gene body and intergenic regions (Figure 5B). Metagene analyses of genes expressed at different levels revealed that the enrichment of phosphorylated RNAPII was associated with gene expression levels in NSN nuclei (Figure 5C). The enrichment of phosphorylated RNAPII at the promoter regions over the gene body regions of highly transcribed genes positively correlated with the open chromatin at the transcription start site (TSS) (Figures 5D and 5E), indicative of active transcription with promoter-proximal pausing^53^. Unexpectedly, different from fertilized eggs^47,54^, phosphorylated RNAPII also accumulated at the 3’ distal intergenic regions, coinciding with the unusual open chromatin at the transcription termination site (TTS) that is absent in SN oocytes (Figures 5D and 5E). Because such enrichment did not result in transcription read-through (Figures 5F and S8C), it was likely due to pausing during transcription termination^55^. Thus, RNAPII degradation specifically clears RNAPII from transcriptionally active chromatin during NSN-to-SN transition.

**Figure 5.**
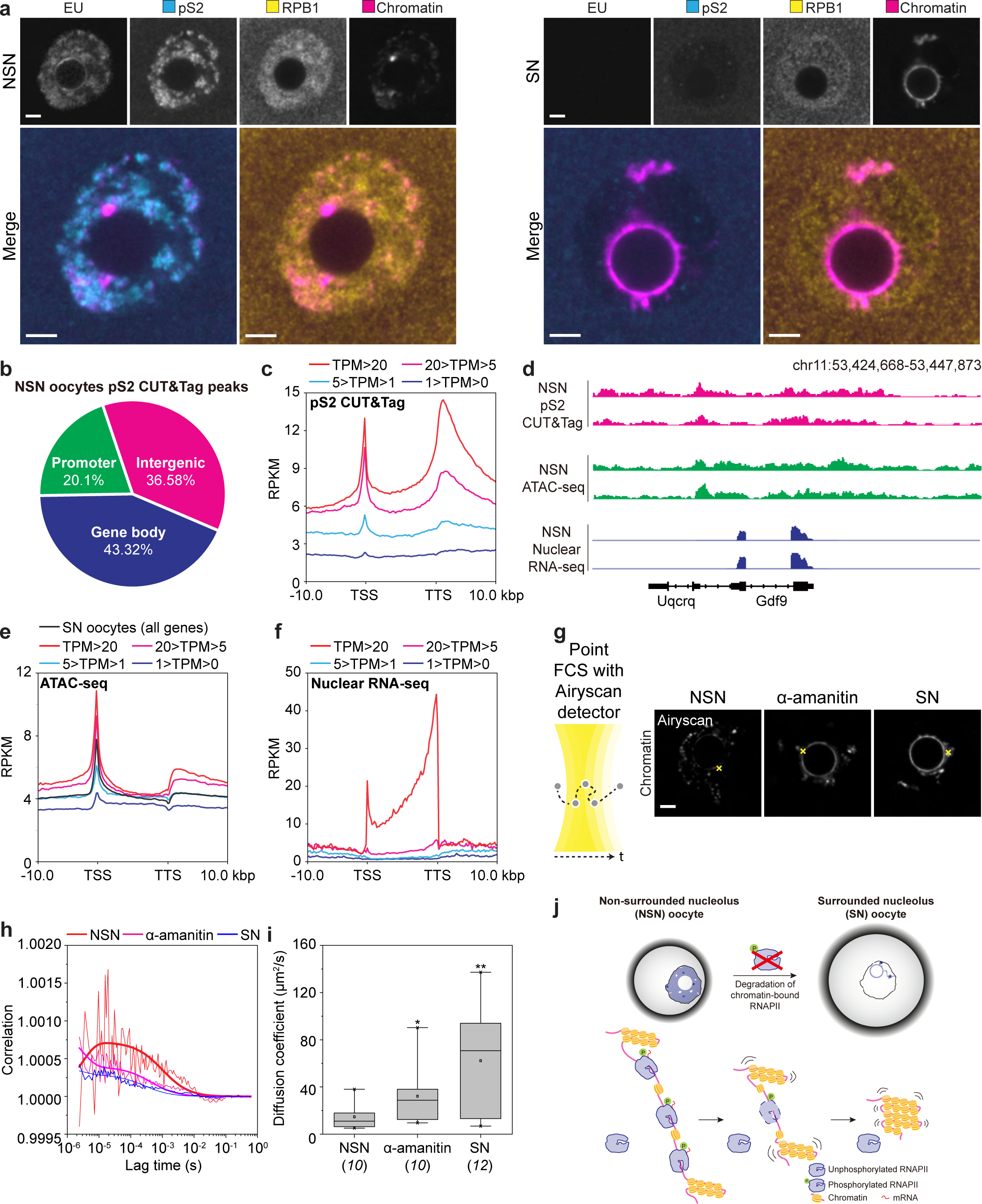
RNAPII degradation globally clears RNAPII from transcriptionally active chromatin and increases chromatin dynamics during NSN-to-SN transition. **(A)** Immunofluorescence images of NSN and SN oocytes. Cyan, pS2; Yellow, RPB1; Magenta, chromatin (Hoechst). **(B)** Pie chart showing the distribution of phosphorylated RNAPII at different genomic regions in NSN oocytes. **(C)** Metagene plot of genes expressed at different levels for phosphorylated RNAPII enrichment. **(D)** IGV browser views showing pS2 CUT&Tag, chromatin accessibility and nuclear RNA-seq signals of a representative gene in NSN oocytes. **(E)** Metagene plot of genes expressed at different levels in NSN oocytes and all genes in SN oocytes for chromatin accessibility. **(F)** Metagene plot of genes expressed at different levels in NSN oocytes for nuclear RNA-seq coverage. **(G)** Point FCS measurement of chromatin mobility in NSN, α-amanitin-treated NSN and SN oocytes. Yellow crosses highlight spots of point FCS measurement in representative oocytes. **(H)** Raw and fitted (in bold) FCS curves for chromatin of a representative NSN, α-amanitin-treated NSN and SN oocytes. **(I)** Quantification of diffusion coefficient of chromatin in NSN, α-amanitin-treated NSN and SN oocytes. **(J)** Mechanistic model for NSN-to-SN transition during mammalian oocyte development. See text for details. *P < 0.05. **P < 0.01. See Materials and methods for box plot specifications. The number of oocytes analyzed is specified in italics. Scale bars are 5 µm.

Studies in other systems reported that chromatin-bound RNAPII could sterically constrain chromatin movement^56,57^, leading us to a model whereby chromatin occupancy by RNAPII restricts chromatin mobility, preventing NSN-to-SN transition from taking place. Consistent with our model, point fluorescence correlation spectroscopy (point FCS) revealed a higher chromatin dynamic in α-amanitin-treated NSN and SN oocytes as compared to NSN oocytes (Figures 5G-5I). To further consolidate our model, we also tested whether NSN-to-SN transition can be induced by reducing the levels of chromatin-bound RNAPII without degrading it. To this end, we treated NSN oocytes with THZ1 and DRB, inhibitors of CDK7 and CDK9 that sequentially phosphorylates RNAPII during the active transcription cycle, respectively. As expected, THZ1 was more effective than DRB in inhibiting transcription and reducing phosphorylated RNAPII levels without affecting total RNAPII levels (Figures S9A-S9D). Importantly, we found that DRB only partially released chromatin-bound RNAPII concomitant with a partial NSN-to-SN transition, whereas THZ1 fully released chromatin-bound RNAPII concomitant with a full NSN-to-SN transition (Figures S9A, S9E and S9F). Thus, RNAPII degradation increases chromatin dynamics, promoting chromatin rearrangement during NSN-to-SN transition.

## Discussion

Our data elucidate a previously unknown principle of chromatin reorganization in mammalian oocytes: RNAPII degradation drives the transition of chromatin from the NSN to SN configuration (Figure 5J). NSN-to-SN transition and transcriptional silencing are two unique features of mammalian oocyte development. It is generally believed that independent mechanisms drive NSN-to-SN transition and transcriptional silencing in oocytes^29,39,58,59^. We challenge this conventional view by showing that RNAPII degradation clears RNAPII from transcriptionally active chromatin and increases chromatin dynamics, simultaneously driving transcriptional silencing and NSN-to-SN transition. This elegant mechanism ensures that transcriptional silencing is tightly coupled with chromatin condensation and rearrangement during NSN-to-SN transition in oocytes.

In this study we systematically examined the changes of many nuclear components during NSN-to-SN transition and directly tested their potential contributions by mimicking their changes using small-molecule inhibitors and overexpression. Many of them are epigenetic marks and factors, which have been implicated in mammalian oocyte development. While modulating many of these factors can alter transcriptional activity and/or chromatin compaction in oocytes, they are not sufficient to induce or reverse NSN-to-SN transition. Nevertheless, we do not exclude the possibility that they help to ensure complete transcriptional silencing and/or chromatin compaction in fully grown oocytes^16,44,59–64^.

Degrading RNAPII using triptolide, α-amanitin or auxin-inducible degron results in little changes in the gross morphology of chromatin in other systems^56,65–70^, including early mammalian embryos^71–76^. Why does RNAPII degradation have more profound impacts on chromatin in mammalian oocytes than in other systems? Recent studies imply that chromatin-bound RNAPII levels increase with and serve as a limiting factor for cell size^77,78^. As growing oocytes expand in size, they continue to load RNAPII into the nucleus and accumulate more phosphorylated RNAPII on chromatin. Indeed, RNAPII levels in NSN oocytes were around 12.5- and 2-fold higher than that in somatic cells or blastomeres, respectively. This may explain the more profound effects on the chromatin in mammalian oocytes upon the clearance of chromatin-bound RNAPII.

Finally, we successfully used scFv for Trim-Away of a non-shuttling nuclear protein. Both fragment antigen-binding (Fab) and its genetically encoded form scFv have been extensively used for imaging endogenous proteins before^79,80^. scFv has advantages of its smaller size and not artificially clustering antigens over the bivalent IgGs^35^, and advantages of higher stability and wider availability over nanobodies^37^. The direct fusion of scFv to t21R also circumvents the need of co-delivering Trim21, opening unprecedented opportunities for using off-the-shelf scFvs like 42B3^36^ to study the functions of nuclear proteins in primary or nondividing cells without the need of any genetic manipulation. Although chemical tools like triptolide and α-amanitin are available, they have pleiotropic effects in different systems and how they induce RNAPII degradation is still largely unclear. The addition of our 42B3-t21R construct to the current toolbox will be invaluable for studying the acute effects of depleting endogenous RNAPII in different mammalian species and cell types.

## Materials and Methods

Extended materials and methods are provided in SI Appendix.

### Husbandry and housing conditions of experimental animals

All mice were maintained in a specific pathogen-free environment at the Animal and Plant Center of the National Institute of Biological Sciences, Beijing according to international animal welfare rules. All experiments with mice compiled with ethical guidelines of the Animal Ethics Committee of the National Institute of Biological Sciences, Beijing.

### Preparation and culture of mouse oocytes and follicles

Oocytes were isolated from ovaries of 8-to 10-week-old FVB/N or C57BL/6J female mice (Charles River Laboratories). GV oocytes were arrested at prophase in homemade phenol red-free M2 supplemented with 250 μM dibutyryl cyclic AMP (dbcAMP) (Sigma-Aldrich or MedCehmExpress) under paraffin oil (ACROS Organics) at 37°C.

Follicles were enzymatically isolated from ovaries of 14-day-old C57BL/6J female mice in L-15 medium (Sigma-Aldrich) supplemented with 5% fetal bovine serum (FBS) (Sigma-Aldrich), 1x GlutaMAX (Thermo Fisher Scientific) and 0.1x penicillin G/streptomycin (Sigma-Aldrich) containing 1 mg/ml collagenase A (Roche) for 15 min at 37°C. Intact follicles of around 100 μm in diameter with a centered oocyte were cultured in MEM alpha with GlutaMAX and nucleosides (Thermo Fisher Scientific) supplemented with 5% FBS, 0.03 μg/ml ovine follicle stimulating hormone (National Hormone and Peptide Program), 1x insulin/transferrin/sodium selenite (Sigma-Aldrich), and 0.1x penicillin G/streptomycin on 12-mm Transwell-COL collagen-coated 0.4-μm pore polytetrafluoroethylene (PTFE) membrane insert (Corning) at 37°C and 5% CO2. Half of the medium surrounding the insert was replaced every 3 days. After 3 to 9 days of culture, in vitro grown oocytes were denuded in M2 with dbcAMP.

### Preparation, culture of mouse MII oocytes and IVF

4-week-old C57BL/6J female mice were superovulated by sequential injection of 0.1 ml CARD HyperOva (Cosmo Bio) and 7.5 IU human chorionic gonadotropin (hCG) (ProSpec) after 48 h. Sperm was collected from caudae epididymides from >12-week-old FVB/N male mice and capacitated for 1 h in 400 μl of homemade phenol red-free HTF. MII oocytes were collected from oviducts 15 h after hCG and briefly cultured in M2. IVF was performed in 100 μl of HTF by adding 10 μl of 3.33×10^7^/ml capacitated sperm for 3 h. Fertilized eggs were then washed in 1:1 HTF:EmbryoMax Advanced KSOM Medium (Sigma-Aldrich) for 30 min and further transferred to EmbryoMax Advanced KSOM Medium for long-term culture. MII oocytes, 1-cell embryos, early and late 2-cell embryos were collected at 0, 7, 22 and 28 h post insemination, respectively.

For in vitro fertilization of GV oocytes, oocytes with a centered nucleus were first treated with water or 100 μM α-amanitin (APExBIO) in M2 with dbcAMP for 7 h and then released into dbcAMP-free M2 for 13.5 h. Before the addition of sperm, a single hole was created in the zona pellucida of the MII oocytes using XYRCOS (Hamilton Thorne). IVF was performed as described above.

### Preparation and culture of human oocytes and follicles

The use of unfertilized human oocytes in this study was approved by the Institutional Review Board (IRB) at the National Institute of Biological Sciences, Beijing under the reference IRB22101001, Reproductive and Genetic Hospital of CITIC-XIANGYA under the reference LL-SC-2017-012 and Beijing Perfect Family Hospital under the reference 2020-09-08. The use of human ovarian cortical tissues in this study was approved by the IRB at the Reproductive and Genetic Hospital of CITIC-XIANGYA under the reference LL-SC-2021-010.

Oocytes were collected from patients who underwent ovarian stimulation for intracytoplasmic sperm injection (ICSI) as part of their assisted reproduction treatment at the Reproductive and Genetic Hospital of CITIC-XIANGYA and Beijing Perfect Family Hospital. Only oocytes that were GV at the time of ICSI and thus unsuitable for the procedure were used in this study. All patients gave informed consent for their surplus oocyte(s) to be used in this study. To maximize the survival and developmental rate of vitrified-thawed GV oocytes^22^, oocytes were vitrified by transferring to 20 μl of G-MOPS PLUS (Vitrolife). 20 μl of G-MOPS PLUS containing 7.5% DMSO and 7.5% ethylene glycol (Sigma-Aldrich) was then added and oocytes were incubated for 3 min at room temperature twice. 240 μl of G-MOPS PLUS containing 7.5% DMSO and 7.5% ethylene glycol were further added and oocytes were incubated for 6 to 10 min at room temperature. Before loading onto Cryotop, oocytes were washed in 300 μl of G-MOPS PLUS containing 15% DMSO, 15% ethylene glycol and 0.5 M D-(+)-trehalose (Sigma-Aldrich) twice. For thawing, Cryotop was inserted into 1 ml of prewarmed G-MOPS PLUS containing 1 M D-(+)-trehalose for 1 min at 37°C. Oocytes were then transferred to 300 μl of G-MOPS PLUS containing 0.5 M D-(+)-trehalose for 3 min at room temperature, 300 μl of G-MOPS PLUS containing 0.25 M D-(+)-trehalose for 5 min at room temperature, and 300 μl of G-MOPS PLUS for 2 min at room temperature. Recovered oocytes were transferred to G-MOPS PLUS at 37°C. Fresh oocytes were directly transferred to and cultured in G-MOPS PLUS at 37°C within 3 to 5 h after retrieval from ovaries. To select for the most immature oocytes, only oocytes that were morphologically normal but did not undergo GV breakdown within 24 h after retrieval from ovaries were used in this study. Follicles were collected from human ovarian cortical tissues retrieved from polycystic ovary syndrome patients who had significantly higher number (≥80) of follicles on ultrasound and required surgical ovarian wedge resection to avoid ovarian hyperstimulation syndrome after having obtained informed consent. Ovarian cortical tissues were cut into 0.5×0.5×1 mm pieces with a Mcllwain Tissue Chooper (Mickle Laboratory Engineering) and digested in MEM alpha with GlutaMAX and nucleosides supplemented with 1x penicillin G/streptomycin (Thermo Fisher Scientific), 0.04 mg/ml Liberase DH (Roche) and 10 IU/ml DNase I (Sigma-Aldrich) for 120 min at 37°C. Digestion was terminated by washing with PBS with 10% BSA twice. Follicles were manually picked and fixed in PBS with 4% paraformaldehyde for 30 min at 37°C before washing in M2.

## Acknowledgments

We are grateful to the patients who participated in this study. We thank the staff from the Transgenic Animal Center & Animal and Plant Center, Imaging Facility and Bioinformatics Center at the National Institute of Biological Sciences, Beijing for technical assistance; the clinicians, nursing team, and embryology team at the clinics for their support of this study; Z. Han and Z. Li for help with human oocytes and mouse embryos; A. Andersen, Q. Liu, and X. Wang for critical comments on the manuscript; M. Schuh for cDNAs and constructs. The research leading to these results was funded by the National Institute of Biological Sciences, Beijing, Ministry of Science and Technology of the People’s Republic of China under a National Key Research and Development Program of China to C.S. (2023YFA1800300) and the National Natural Science Foundation of China (NSFC) under a grant to J.G. (82371667). C.S is also a recipient of the Excellent Young Scientists Fund Program from the NSFC and the DAMO Academy Young Fellowship from the Alibaba DAMO Academy. The authors apologize to colleagues whose work could not be cited owing to length limitations.

## Author contributions

J.W. and C.S. conceived the study, designed experiments and methods for data analysis; J.W. and C.S. performed all experiments and analyzed the data with the following exceptions: W.L. performed bioinformatics analysis of all next generation sequencing data. J.W. and C.S. wrote the manuscript and prepared the figures with input from all authors; J.G. and G.L. supervised the collection of human ovarian cortical tissues and vitrification of human oocytes at the Reproductive and Genetic Hospital of CITIC-XIANGYA; X.X. supervised the collection and vitrification of human oocytes at the Beijing Perfect Family Hospital; B.L. supervised the bioinformatics analyses; and C.S. supervised the entire study.

## Supplemental Information

## Materials and Methods

Figures S1-S9 and Table S1

Videos S1 to S8

## Materials and Methods

### Drug treatment

All drugs except 1,6-hexanediol were prepared in water for embryo transfer or DMSO (Sigma-Aldrich). Oocytes were treated with 20 μM 8-AA (MedChemExpress), 20 μM A-485 (MedChemExpress), 100 μM α-amanitin, 100 nM ActD (MedChemExpress), 6 μM BRM/BRG1i (MedChemExpress), 10 μM bleomycin (MedChemExpress), 10 μg/ml CHX (Sigma-Aldrich), 100 μM DRB (MedChemExpress), 10 μM DZNep (MedChemExpress), 25 μg/ml etoposide (MedChemExpress), 100 μM GSK-J4 (MedChemExpress), 500 μM hinokiflavone (MedChemExpress), 10 μM quarfloxin (MedChemExpress), 2.5 μM THZ1 (MedChemExpress), 10 μM triptolide (MedChemExpress), 100 nM TSA (MedChemExpress) or 10 μM 7-DAA (MedChemExpress) in M2 with dbcAMP for 7 h. For 1,6-hexanediol treatment, oocytes were treated with 15% 1,6-hexanediol (Sigma-Aldrich) in M2 with dbcAMP for 10 min. For CHX treatment, oocytes were treated with 10 μg/ml CHX in M2 with dbcAMP for 4 to 24 h. For CB-5083, MG132 and flavorpiridol treatments, cultured mouse ovarian follicles were treated with 5 μM CB-5083 (MedChemExpress), 25 μM MG132 (MedChemExpress) or 5 μM flavorpiridol (MedChemExpress) on D14+7 for 48 h.

### Expression constructs and mRNA synthesis

To generate constructs for mRNA synthesis, we fused previously published coding sequences with eGFP, mClover3, P2A, NLS(NP), t21R and/or V5 and subcloned them into pGEMHE to obtain 42B3-mClover3, 42B3-mClover3(11)-P2A-NLS-mClover3(1-10) (42B3-split mClover3), 42B3-t21R, mClover3-CTCF, DNMT1-V5, DNMT3A-V5, DNMT3B-V5, DNMT3L-V5, NLS-eGFP-hnRNPU, HP1β, NLS-eGFP-MATR3, mClover3-RPB1, NLS-mClover3-RPB1, TET1-V5, TET2-V5 and TET3-V5. All mRNAs were synthesized and quantified as previously described^22^.

### Microinjection of mRNAs

Mouse oocytes were microinjected with 3.5 to 14 pl of mRNAs as previously described^22^. *42B3-mClover3* mRNA was microinjected at a needle concentration (final concentration in the microinjection needle) of 560 ng/μl, *42B3-split mClover3* mRNA at 560 ng/μl, *42B3-t21R* mRNA at 2965.9 ng/μl, *mClover3-CTCF* mRNA at 3040 ng/μl, *DNMT1-V5* mRNA at 572.4 ng/μl, *DNMT3A-V5* mRNA at 701 ng/μl, *DNMT3B-V5* mRNA at 752.2 ng/μl, *DNMT3L-V5* mRNA at 1057 ng/μl, *NLS-eGFP-hnRNPU* mRNA at 2251.2 ng/μl, *HP1β* mRNA at 952.5 ng/μl, *NLS-eGFP-MATR3* mRNA at 1148.1 ng/μl, *mClover3-RPB1* mRNA at 2185.5 ng/μl, *NLS-mClover3-RPB1* mRNA at 2424.6 ng/μl, *TET1-V5* mRNA at 801.9 ng/μl, *TET2-V5* mRNA at 528.9 ng/μl and *TET3-V5* mRNA at 469.3 ng/μl. Oocytes were allowed to express the mRNAs for 3 to 4 h.

Human oocytes were microinjected with 14 pl of *42B3-t21R* mRNA at a needle concentration of 2965.9 ng/μl as previously described^22^.

### Staining of GV oocytes for visualizing chromatin configuration and live imaging

SPY555-DNA (Spirochrome) and 5-SiR-Hoechst^22^ was reconstituted in DMSO. For visualizing chromatin configuration, oocytes were stained with SPY555-DNA or 5-SiR-Hoechst for 30 min at 1:500 dilution and 10 μM, respectively. For live imaging, SPY555-DNA or 5-SiR-Hoechst was present throughout imaging at 1:1000 dilution and 50 nM, respectively.

### Immunofluorescence staining

Oocytes were fixed in 100 mM HEPES (pH 7.0, titrated with KOH), 50 mM EGTA (pH 7.0, titrated with KOH), 10 mM MgSO_4_, 2% methanol-free formaldehyde, and 0.5% triton X-100 for 15 to 60 min at 37°C and washed in phosphate-buffered saline (PBS). For EU and L-AHA labelling, oocytes were pulsed with 1 mM EU (Thermo Fisher Scientific) or 100 μM L-AHA (Click Chemistry Tools) in M2 for 2 h at 37°C before fixation and visualized using Click-iT RNA Alexa Fluor 488 Imaging Kit (Thermo Fisher Scientific) or AFDye 568 Azide (Click Chemistry Tools) or Click-iT HPG Alexa Fluor 488 Protein Synthesis Assay Kit (Thermo Fisher Scientific). For 5mC and 5hmC staining, fixed oocytes were treated with 4 M HCl for 10 min and neutralized with 100 mM Tris-HCl (pH 7.5) for 10 min. All antibody incubations and washings were performed in PBS with 0.5% triton X-100 (PBT) and 5% BSA (PBT-BSA) at 10 μg/ml overnight at 4°C (for primary antibodies) and at 20 μg/ml for 1 h at room temperature (for secondary antibodies). Primary antibodies used were mouse anti-B23 (Santa Cruz Biotechnology #sc-56622), rabbit anti-CTCF (Sigma-Aldrich #07-729-25UL), mouse anti-5mC (Active Motif #39649), rabbit anti-5hmC (Active Motif #39792), rabbit anti-FBL (Abcam #ab166630), goat anti-GFP (Rockland Immunochemicals #600-101-215), mouse anti-γH2A.X (Sigma-Aldrich #05-636), rabbit anti-H3ac (Active Motif #39140), rabbit anti-H4ac (Sigma-Aldrich #06-598), rabbit anti-hnRNPU (Huabio #ET7107-10), rabbit anti-HP1α (Huabio #ET1602-8), mouse anti-HP1β (Sigma-Aldrich #MAB3448), rabbit anti-HP1γ (Huabio #ET1706-02), rabbit anti-MATR3 (Huabio #ET7106-95), rabbit anti-HMe2 #1 (Abcam #ab7315), rabbit anti-HMe2 #2 (Abclonal #A5870), rabbit anti-HMe2 #3 (Abclonal #A18296), rabbit anti-HMe3 #1 (Thermo Fisher Scientific #PA5-116816), rabbit anti-HMe3 #2 (Abclonal #A20145), rabbit anti-NuMA (Abcam #ab97585), rabbit anti-POLR1A (CST #24799), rabbit anti-RPB1 (CST #149158S), mouse anti-CTD pY1 (Active Motif #91220), rat anti-CTD pS2 (Active Motif #61999), rabbit anti-CTD pT4 (Abclonal #AP1383), rat anti-CTD pS5 (Active Motif #67102), rat anti-CTD pS7 (Active Motif #61704), mouse anti-SRSF2 (Sigma-Aldrich #S4045), rabbit anti-SMARCA2 (Abclonal #A23291), rabbit anti-SMARCA4 (Abclonal #A19556), rabbit anti-SMC3 (Abcam #Ab128919) and rabbit anti-SMC4 (Sigma-Aldrich #HPA029449). Secondary antibodies used were Alexa Fluor 488-, 594-, or 647-conjugated AffiniPure Fab Fragment anti-goat IgG, anti-rabbit IgG, and anti-rat IgG (Jackson ImmunoResearch Europe), Alexa Fluor 568-conjugated Nano-Secondary anti-mouse IgG1, IgG2a and IgG2b (Proteintech). DNA was stained with Hoechst 33342 (Molecular Probes).

### polyA RNA-FISH

Oocytes were fixed in 100 mM HEPES, 50 mM EGTA, 10 mM MgSO_4_, 2% methanol-free formaldehyde, and 0.5% triton X-100 for 15 to 60 min at 37°C and washed in PBT. Oocytes were then washed with Wash Buffer A (Biosearch Technologies) containing 10% formamide and incubated with 200 nM Alexa Fluor 647-conjugated oligo dT30 (IDT) in hybridization buffer (Biosearch Technologies) supplemented with 10% formamide overnight at 37°C. After probe incubation, oocytes were washed with Wash Buffer A for 30 min at 37°C and stained with Hoechst 33342 in Wash Buffer A for another 30 min at 37°C. Oocytes were further washed with Wash Buffer A and Wash Buffer B (Biosearch Technologies) for 30 min at 37°C and room temperature, respectively, before imaging.

### RIBOmap

Oocytes were fixed in 100 mM HEPES, 50 mM EGTA, 10 mM MgSO_4_ and 2% methanol-free formaldehyde for 15 to 60 min at 37°C and washed in PBT. Oocytes were then washed three times with hybridization buffer consisting of 2x SSC, 10% formamide, 0.1% triton X-100, 0.1 mg/ml yeast tRNA (Thermo Fisher Scientific) and 0.2 U/μl RiboLock RNase Inhibitor (Thermo Fisher Scientific). Hybridization solution was prepared by diluting all probes (Genewiz), including 5 splints (5’-ACAAAATAGAACCGCGGTCCTATTCAA AAA AAA AAA AAA AAA AAA AAA AAA AAA AAA AAA AAA AAA AAA AAA AAA TAT CTT TAG T*G*T* /3InvdT/-3’; 5’-CATCGTTTATGGTCGGAACTACGACAA AAA AAA AAA AAA AAA AAA AAA AAA AAA AAA AAA AAA AAA AAA AAA AAA TAT CTT TAG T*G*T* /3InvdT/-3’; 5’-AGGTTTCCCGTGTTGAGTCAAATTAA AAA AAA AAA AAA AAA AAA AAA AAA AAA AAA AAA AAA AAA AAA AAA AAA TAT CTT TAG T*G*T* /3InvdT/-3’; 5’-TGTTATTGCTCAATCTCGGGTGGCTAA AAA AAA AAA AAA AAA AAA AAA AAA AAA AAA AAA AAA AAA AAA AAA AAA TAT CTT TAG T*G*T* /3InvdT/-3’; 5’-AGATAGTCAAGTTCGACCGTCTTCTAA AAA AAA AAA AAA AAA AAA AAA AAA AAA AAA AAA AAA AAA AAA AAA AAA TAT CTT TAG T*G*T* /3InvdT/-3’), 5 *Rpb1*-specific padlocks (5’-/5Phos/AAGATA AACATCGTAGACTA GCTACAAAAAGCCTGCGCC TCAGGTCAT ACACTA-3’; 5’-/5Phos/AAGATA AACATCGTAGACTA TGTCAATATTGAAGGGTGTGACAATCT TCAGGTCAT ACACTA-3’; 5’-/5Phos/AAGATA AACATCGTAGACTA TCTATTACATCTTGTTTAGCTTTCTTAATAGTGT TCAGGTCAT ACACTA-3’; 5’-/5Phos/AAGATA AACATCGTAGACTA GTGAAGATCTCCACAATATCATTGGACG TCAGGTCAT ACACTA-3’; 5’-/5Phos/AAGATA AACATCGTAGACTA TGGTGAGGGGATGTATGGGC TCAGGTCATACACTA-3’) and 4 *Rpb1*-specific primers (5’-CTGCGCAGGCGCAAACC GTGTCTACGATG-3’; 5’-CCCCTGCGTACTAATTCCTGAAG GTGTCTACGATG-3’; 5’-CTCATTGTTATGAGCCTTCTCAATGA GTGTCTACGATG-3’; 5’-CACAGCCTCAATGCCCAGTA GTGTCTACGATG-3’; 5’-AGCTGGGAGACATAGCACCAGTGTCTACGATG-3’), to a final concentration of 50 nM with hybridization buffer. Oocytes were incubated with hybridization solution for 4 to 6 h at 40°C. After hybridization, oocytes were washed twice with PBS with 0.1% triton X-100 and 0.1 U/μl RiboLock RNase Inhibitor (PBSTR) and once with 4x SSC diluted with PBSTR. All washes were performed for 10 min at 37°C. Oocytes were then rinsed with PBSTR and incubated for 1.5 h at room temperature with ligation solution consisting of 0.25 Weiss U/μl T4 DNA ligase (Thermo Fisher Scientific), 1x T4 DNA ligase buffer, 0.5 mg/ml BSA and 0.4 U/μl RiboLock RNase Inhibitor. After two washes with PBSTR for 5 min each, oocytes were incubated with amplification-labeling mixture consisting of 0.5 U/μl phi29 DNA polymerase (NEB), 1x phi29 DNA polymerase reaction buffer, 250 μM dNTP (NEB), 0.5 mg/ml BSA, 0.4 U/μl RiboLock RNase Inhibitor and 1 μM labeling probe (5’-/5Cy3/CATACACTAAAGATAACAT-3’) (Genewiz) for 2 h at 30°C. Oocytes were subsequently washed twice with PBSTR with 0.05 mg/ml BSA for 5 min each at room temperature before imaging.

### Confocal and light-sheet microscopy

For confocal imaging, fixed oocytes were imaged in 2 μl of medium (for live oocytes) or PBS with 1% polyvinylpyrrolidone (PVP) and 0.5 mg/ml BSA (for fixed oocytes) under paraffin oil in a 35-mm dish with a no. 1.5 coverslip. Images were acquired with LSM 900 (Zeiss), LSM 980 (Zeiss) and Stellaris 5 (Leica) confocal laser scanning microscope equipped with a 40x C-Apochromat 1.2 NA water-immersion objective (Zeiss) or a 63x Plan-Apochromat 1.4 NA oil-immersion objective (Leica). mClover3 and Alexa Fluor 488 were excited with a 488-nm laser line and detected at 493 to 556 nm. Alexa Fluor 568 and Alexa Fluor 594 were excited with a 561-nm laser line and detected at 566 to 635 nm. Alexa Fluor 647 and 5-SiR-Hoechst were excited with a 633-or 640-nm laser line and detected at 645 to 700 nm.

For light-sheet imaging, oocytes were imaged in 0.5 ml of medium in four different compartments of a multiwell sample holder (Viventis Microscopy Sàrl). Dual-view images were acquired with LS2 Live (Viventis Microscopy Sàrl) equipped with an environmental incubator box and two 25x 1.1 NA water-dipping objectives (Nikon). SPY555 was excited with a 561-nm laser line and detected with a 523/20 nm – 610/25 nm dual band-pass filter. Dual-view images were processed with a fusion script (Viventis Microscopy Sàrl) after acquisition.

Images of the control and experimental groups were acquired under identical imaging conditions on the same microscope. For some images, shot noise was reduced with a Gaussian filter. Care was taken that the imaging conditions (laser power, pixel-dwell time, detector gain, and exposure time) did not cause phototoxicity (for live imaging), photobleaching and saturation.

### Image quantification

To reproducibly segment chromatin (labeled with Hoechst 33342 or SPY555-DNA) or other nuclear structures (labeled with antibodies against different nuclear components), the Machine Learning Segmenter was used for automatic segmentations in Arivis (Zeiss). For each experiment, 3 to 5 z-planes from 5 to 10 cells or timepoints were manually annotated for foreground and background pixels and subjected to model training under the Fluorescence Fast mode. Suitable smoothing and threshold values were selected to refine the segmentation results before batch applications. Total fluorescence intensity, volume and surface area of the segmented objects were exported into Excel (Microsoft) and OriginPro (OriginLab) for further processing.

### Point FCS

For analyses of chromatin mobility, point FCS was performed on oocytes stained with 5-SiR-Hoehcst using the Dynamics Profiler and Airyscan modules (Zeiss) on LSM 980 equipped with an environmental incubator box (Peacon). Spot regions of interests (ROIs) were marked and measured for fluorescence fluctuations using the 640-nm laser line for 10 s. Data were first processed and then fitted to obtain the diffusion coefficient. Specific parameters used were: 150 for detrend and 1 Component 3D for fit model.

### Immunoblotting

40 mouse oocytes or embryos (per lane) were extensively washed in protein-free medium and snap-frozen in 1 μl of protein-free medium in liquid nitrogen. Before thawing, 7 μl of water and 4 μl of 4x NuPAGE LDS sample buffer (Thermo Fisher Scientific) with 100 mM dithiothreitol (DTT) was added. Samples were then thawed at 37°C and snap-frozen in liquid nitrogen twice more before being heated for 5 min at 99°C. For λPP treatment, oocytes were lyzed in 8 μl of buffer (1X NEBuffer for protein metallophosphatase and 1 mM MnCl_2_) by snap-freezing in liquid nitrogen and thawing at 37°C three times, and incubated for 30 min at 30°C after adding 0.16 μl of 400 U/μl λPP. Samples were resolved on a 15-well NuPAGE 8 to 16% Tris-Glycine Plus WedgeWell Gel of 1.0 mm thickness (Thermo Fisher Scientific) with Tris-Glycine SDS Running Buffer (Thermo Fisher Scientific or Beyotime). Proteins were transferred onto a methanol-activated 0.45-μm polyvinylidene difluoride (PVDF) membrane (Thermo Fisher Scientific) with SDS-free Towbin buffer at 200 mA for 2 h on ice. Blots were stained with No-Stain Protein Labeling Reagent (Thermo Fisher Scientific) before blocking. Blocking and antibody incubations were performed in Tris-buffered saline (TBS) with 5% skim milk and 0.1% tween-20. Primary antibodies used were rabbit anti-RPB1 (CST #149158S), mouse anti-CTD pY1 (Active Motif #91220), rat anti-CTD pS2 (Active Motif #61699), rabbit anti-CTD pT4 (CST #26319S), mouse anti-CTD pS5 (4H8) (Santa Cruz sc-47701) and rat anti-TUBA (Bio-Rad #MCA78G). Secondary antibodies used were horseradish peroxidase (HRP)-conjugated goat anti-rabbit IgG (CST #7074P2), goat anti-rat IgG (CST #7077S) and horse anti-mouse IgG (CST #7076P2). Blots were developed with SuperSignal West Femto Maximum Sensitivity Substrate (Thermo Fisher Scientific) and documented with Amersham ImageQuant 500 (Cytiva). Care was taken that the exposure time did not cause saturation.

### RT-qPCR

40 mouse oocytes (per sample) were extensively washed in protein-free medium and snap-frozen in 1 μl of protein-free medium in liquid nitrogen. Total RNA was extracted with TRIzol LS Reagent (Thermo Fisher Scientific) in Phasemaker Tube (Thermo Fisher Scientific) and treated with TURBO DNA-free Kit (Thermo Fisher Scientific). Reverse transcription (RT) was performed using SuperScript IV First-Strand Synthesis System (Thermo Fisher Scientific), and quantitative PCR (qPCR) was performed using SsoAdvanced Universal SYBR Green Supermix (Bio-Rad) on CFX Opus 96 Real-Time PCR System (Bio-Rad). Endogenous *Rpl13a* was used as an internal control to calculate relative transcript level. For amplifying *Rpl13a*, the following primers were used: 5’-GAGGTCGGGTGGAAGTACCA-3’ and 5’-TGCATCTTGGCCTTTTCCTT-3’. For amplifying *Rpb1*, the following primers were used: 5’-ACATGTGCAGGAAACATGAC-3’ and 5’-CTTCGGGTTATTAGAATCTACAAGC-3’.

### Poly(A) Tail-Length Assay

40 mouse oocytes (per sample) were extensively washed in protein-free medium and snap-frozen in 1 μl of protein-free medium in liquid nitrogen. Total RNA was extracted with TRIzol LS Reagent in Phasemaker Tube and treated with TURBO DNA-free Kit. G/I tailing, RT and end-point PCR were performed according to the manual of the Poly(A) Tail-Length Assay Kit (Thermo Fisher Scientific). The poly(A) start site (PAS) fragment was obtained by PCR using the *Rpb1*-specific forward primer (5’-TTGGTGCCTGCTCTGG-3’) and the *Rpb1*-specific reverse primer (5’-TGGTCAAAATTAGTAAACTTTATTTCAATTTCAAAAAATAACAAA-3’) located immediately upstream of the PAS. The tail fragment was obtained by PCR using the same *Rpb1*-speicfic forward primer and a 35 nt universal reversal primer provided in the kit. The PCR products were resolved on a 2.5% agarose gel, and the poly(A) tail length was calculated by (size of tail fragment) – (size of PAS fragment) – 35.

### Hi-C, data processing and analysis

Chromatin digestion was performed as previously described^31^ after minor modifications. 100 mouse oocytes (per sample) were fixed in PBS with 0.05% PVP and 1% methanol-free formaldehyde for 10 min at RT and 2.5 M glycine was added to a final concentration of 0.2 M to quench for 5 min at RT. Oocytes were transferred to 100 μl of PBS with 0.05% PVP and captured by adding 10 μl of washed ConA Beads Pro (Vazyme) for 10 min at RT. Oocytes were lysed in 12.5 μl of Hi-C lysis buffer [10 mM Trim-HCl (pH 8.0), 10 mM NaCl, 0.5 % NP-40] supplemented with cOmplete, Mini, EDTA-free Protease Inhibitor Cocktail (Roche) for 15 min on ice. Supernatants were discarded with 2.25 μl remaining, and 0.25 μl of 5% SDS was added before incubating for 10 min at 62°C. 7.25 μl of nuclease-free water and 1.25 μl of 10% Triton X-100 were further added to quench for 15 min at 37°C, and 1.25 μl of 10x NEBuffer 2 (NEB) and 5 U of MboI (NEB) were added to digest chromatin overnight at 37°C. The next day samples were incubated for 20 min at 62°C to inactivate MboI and then cooled to RT. 2.5 μl of fill-in master mix [7.5 μl of 0.4 mM biotin-14-dATP (Active Motif), 0.3 μl of 10 mM dCTP, 0.3 μl of 10 mM dGTP, 0.3 μl of 10 mM dTTP and 1.6 μl of 5 U/μl of DNA polymerase I, Large (Klenow) Fragment (NEB)] was added to fill in restriction fragment overhangs and mark DNA ends with biotin, and samples were incubated for 90 min at 37°C. Ligation master mix [33.2 μl of nuclease-free water, 11 μl of 10x T4 DNA Ligase buffer (NEB), 5 μl of 10% Triton X-100, 0.6 μl of 100x BSA and 0.5 μl of 2000 U/μl T4 DNA Ligase (NEB)] was further added, and samples were incubated for 4 h at RT. After removing supernatants, beads were resuspended with 15.5 μl of nuclease-free water, 1.75 μl of Lambda Exonuclease Buffer (NEB), 0.15 μl of Lambda Exonuclease (NEB) and 0.15 μl of Exonuclease I (NEB)] and incubated for 1 h at 37°C to remove unligated ends. To release DNAs, 42.5 μl of nuclease-free water, 6 μl of 10% SDS and 2.5 μl of 20 mg/ml Proteinase K (Thermo Fisher Scientific) were added, and samples were incubated for 30 min at 55°C. To reverse crosslinks, 6.5 μl of 5 M NaCl was added, and samples were incubated overnight at 68°C. The next day DNAs were extracted with phenol-chloroform in Phasemaker Tube, precipitated with ethanol and resuspended in 5.5 μl of nuclease-free water. Library preparation was performed using the TruePrep DNA Library Prep Kit V2 for Illumina (Vazyme) after minor modifications. DNA fragmentation mix was scaled down to 10 μl. After tagmentation, biotin pull-down process was begun by washing 12.5 μl of 10 mg/ml Sera-Mag SpeedBeads Neutravidin-Coated Magnetic Particles (Cytiva) or Dynabeads MyOne Streptavidin C1 (Thermo Fisher Scientific) with 40 μl of 1x TWB [5 mM Tris-HCl (pH 7.5), 1 M NaCl, 0.5 mM EDTA and 0.05% tween-20]. After resuspension in 25 μl of 2x BB [10 mM Tris-HCl (pH 7.5), 2 M NaCl and 1 mM EDTA], beads were added to samples and incubated for 15 min at RT. Before PCR, beads were washed with 40 μl of 1x TWB twice and resuspended in 29 μl of nuclease-free water. The purified libraries were quantified by Qubit 1X dsDNA HS Assay Kit (Thermo Fisher Scientific) and qPCR, analyzed by a Fragment Analyzer System (Agilent) to confirm successful amplification and quality, and sequenced using Novaseq 6000 Sequencing System (Illumina).

Reads were processed using HiC-Pro (v3.1.0) with default parameters in the HiC-Pro configuration file to obtain valid contacts. The “.bed” file of restriction fragments from MboI digestion was generated using the “digest_genome” Python script from HiC-Pro. Reads were initially mapped to mm10 (GRCm38.p6) reference genome, and unmapped reads were subsequently cut at restriction sites and re-mapped. After removing PCR duplicates and reads lacking restriction sites, contact matrix files were generated from the final set of valid contacts, which were then normalized by the ICE method. For visualization, the “hicpro2juicbox” shell script was employed to convert valid contacts into “.hic” files, which were then imported into Juicebox (v.2.20). For calculation of contact probability within chromosomes, valid contacts were converted into “.cool” files using Cooler (v0.9.3) and Cooltools (v0.6.1), and the resulting contact probability curves were smoothed and aggregated for plotting.

### WGBS, data processing and analysis

30 mouse oocytes (per sample) were lyzed in 20 μl of lysis buffer consisting of 1x M-Digestion Buffer and 1 mg/ml Proteinase K (Zymo Research) for 20 min at 50°C. Bisulfite conversion and library preparation were performed using the Pico Methyl-Seq Library Prep Kit (Zymo Research) after minor modifications. In the amplification with PrepAmp Primer step, PrepAmp primer was diluted from 40 μM to 20 μM. In the library amplification step, 10 cycles were used. The purified libraries were quantified by Qubit 1X dsDNA HS Assay Kit and qPCR, analyzed by a Fragment Analyzer System to confirm successful amplification and quality, and sequenced using Novaseq 6000 Sequencing System.

Reads were processed using TrimGalore (v0.6.7) to remove adaptor and low-quality reads with parameters “-q 20, -e 0.1, --length 20, --stringency 1”, and then trimmed reads were mapped to mm10 (GRCm38.p6) reference genome using Bsmap (v2.90) with default parameters. Methylation level of each CpG site was estimated using the function “mcall” of Moabs (v1.3.9.6). Methylation level of each CpG site from multiple replicates was determined using total methylated read count across replicates versus total read count across replicates, and CpG sites with less than 3 reads were discarded.

### ATAC-seq, data processing and analysis

Library preparation was performed as previously described^49^ after minor modifications. 50 mouse oocytes (per sample) were treated with acidic Tyrode’s solution (Sigma-Aldrich) to remove the zona pellucida and transferred into 500 μl of ATAC-Resuspension Buffer [10 mM Tris-HCl (pH 7.4), 10 mM NaCl and 3 mM MgCl2] containing 0.1% NP-40, 0.1% tween-20 and 0.01% digitonin to lyze for nuclei using a glass capillary with <50 μm opening. The purified libraries were quantified by Qubit 1X dsDNA HS Assay Kit and qPCR, analyzed by a Fragment Analyzer System to confirm successful amplification and quality, and sequenced using Novaseq 6000 Sequencing System.

Reads were processed using TrimGalore (v0.6,7) to remove adaptor and low-quality reads with parameters “-q 20, -e 0.1, --length 20, --stringency 1”, and then trimmed reads were mapped to mm10 (GRCm38.p6) reference genome using Bowtie2 (v2.4.2) with parameters “-L 25, -X 2000, --no-mixed --no-discordant”. Unmapped reads and low mapping quality reads (Q<30) were discarded, and PCR duplicates were removed using the function “MarkDuplicates” from Picard tools (v2.27.2). ATAC-seq bigwig tracks were generated using the function “bamcoverage” from Deeptools (v3.5.1) with parameters “-binSize 50-normSize 2494787188-exactScaling - extendReads-normalizeUsing RPKM”. ATAC signal counts in each genomic region were computed using the function “computeMatrix” from Deeptools (v3.5.1) with parameters “-bs 200, --upstream 10000, --downstream 10000”.

### Fractionated RNA-seq, data processing and analysis

Karyoplast and cytoplast were separated by micromanipulation as previously described^9^. 5 μl of Dynabeads MyOne Streptavidin C1 was washed with 20 μl of Dynabead solution A (50 mM NaCl and 0.1 M NaOH) twice and 20 μl of Dynabead solution B (100 mM NaCl). Dynabeads were then resuspended in 5 μl of 2x B&W buffer [10 mM Tris-HCl (pH 7.5), 2 M NaCl and 1 mM EDTA] and 5 μl of 100 μM biotinylated oligo-dT30VN (Sangon Biotech) was added for 20 min at room temperature. Dynabeads were further washed with 20 μl of 1x B&W buffer four times and resuspended in 10 μl of nuclease-free water. 25 karyoplasts or 3 cytoplasts (per sample) were lyzed in 5 μl of lysis buffer [10 mM Tris-HCl (pH 7.5-8.0), 4% NP-40 and 0.2% SDS], and 0.24 μl of 5 M NaCl and 1 μl of Dynabeads were added for 20 min at room temperature. The supernatant was collected as the polyA-ve fraction, and 1 μl of 10^5^-diluted ERCC RNA Spike-In Mix (Thermo Fisher Scientific) and 6.6 μl of nuclease-free water were added. Dynabeads were resuspended in 5 μl of lysis buffer as the polyA+ve fraction, and 1 μl of 10^5^-diluted ERCC RNA Spike-In Mix and 7.6 μl of nuclease-free water were added. Library preparation was performed using the Ovation SoLo RNA-Seq System (Tecan) after minor modifications. In the primer annealing step, 1 μl of 50 μM random hexamer primer (Thermo Fisher Scientific) was added instead of 2.6 μl of First Strand Primer Mix. The purified libraries were quantified by Qubit 1X dsDNA HS Assay Kit and qPCR, analyzed by a Fragment Analyzer System to confirm successful amplification and quality, and sequenced using Novaseq 6000 Sequencing System.

Reads were processed using TrimGalore (v0.6.7) to remove adaptor and low-quality reads with parameters “-q 20, -e 0.1, --length 20, --stringency 1”, and then trimmed reads were mapped to mm10 (GRCm38.p6) reference genome plus ERCC.fasta using STAR (v2.7.10a) with default parameters. Multi-mapping reads were filtered using Samtools (v1.17), and PCR duplicates were removed using the function “MarkDuplicates” from Picard tools (v.2.27.2). Deduplicated reads were mapped to genes using HTseq (v2.0.3) as unstranded samples with default parameters. A gene/ERCC count was considered valid when present in at least five reads in at least two samples. Differentially expressed genes between groups of samples were identified using DESeq2 (v1.38.0). Gene counts of each sample were normalized by defining ERCC genes as the “controlGenes” when estimating the “sizeFactors” with DESeq2.

### Ribo-seq, data processing and analysis

Library preparation was performed as previously described^50^ after minor modifications. 50 mouse oocytes (per sample) were lyzed in 150 μl of ice-cold lysis buffer [20 mM Tris-HCl (pH 8.0), 150 mM NaCl, 5 mM MgCl2, 1% triton X-100 and 1 mM DTT] for 5 min on ice followed by snap-freezing in liquid nitrogen and thawing at 37°C three times. After spinning at 21300 xg for 5 min at 4°C, the supernatant was digested with 0.25 μl of 2 U/μl TURBO DNase (Thermo Fisher Scientific) and 0.2 μl of 1:10-diluted 20 U/μl RNase T1/0.5 U/μl RNase A cocktail (Thermo Fisher Scientific) for 30 min at 37°C. 5 μl of 20 U/μl SUPERaseIn RNase inhibitor (Thermo Fisher Scientific) and 95 μl of ice-cold lysis buffer were further added, and the mixture was extracted using TRIzol LS Reagent in Phasemaker Tube. Purified RNAs were resuspended in 5 μl of PNK solution [5 U T4 PNK (NEB) in 1X T4 PNK reaction buffer] and incubated for 10 min at 37°C. After pre-incubation, ATP was added to a final concentration of 10 mM and the mixture was incubated for another 30 min at 37°C before adding 5.5 μl of Gel Loading Buffer II (Thermo Fisher Scientific). Ribosome-protected fragments (RPFs) corresponding to the 25 to 35 nt band were separated using 6% TBE-Urea Gel (Thermo Fisher Scientific) and subjected to library preparation using NEXTFLEX Small RNA Seq Kit v3 (Perkin Elmer). The purified libraries were quantified by Qubit 1X dsDNA HS Assay Kit and qPCR, analyzed by a Fragment Analyzer System to confirm successful amplification and quality, and sequenced using Novaseq 6000 Sequencing System.

Reads were processed using TrimGalore (v0.6.7) to remove adaptor and low-quality reads with parameters “-q 20, --phred33, --stringency 3, --length 20, --max_length 40”. As the NEXTFLEX Small RNA Seq Kit v3 randomly added 4 bases at the 5’ and 3’ end of each read, we clipped these random bases when trimming reads. Reads mapped to rRNA were removed using Bowtie (v1.3.1) with parameters “-v 1, -a, --best, --strata, -q, --phred33-quals”. Remaining reads were mapped to mm10 (GRCm38.p6) reference genome using STAR (v2.7.10a) with parameters “--outFilterType BySJout, -outFilterMismatchNmax 2, --outFilterMultimapNmax 1, - outFilterMatchNmin 16, -alignEndsType EndToEnd, -outSAMattrIHstart 0”. Differentially translated mRNAs were identified using DESeq2 (v1.38.0).

### CUT&Tag, data processing and analysis

Library preparation was performed as previously described^49^ after minor modifications. 50 to 80 mouse oocytes (per sample) were pre-extracted with ice-cold extraction buffer [25 mM HEPES (pH 7.4), 50 mM NaCl, 3 mM MgCl2, 300 mM sucrose and 0.5% triton X-100] for 10 min on ice and washed three times through ice-cold extraction buffer without triton X-100. Pre-extracted oocytes were directly washed with antibody buffer and incubated with primary antibody without crosslinking. Primary antibodies used were mouse IgG (Beyotime #A7028), mouse anti-CTD pS5 (4H8) (Santa Cruz #sc-47701), rat IgG (Beyotime #A7031), rat anti-CTD pS2 (Active motif #61699). Secondary antibodies used were rabbit anti-mouse (Sigma-Aldrich #06-371) and rabbit anti-rat (Sigma-Aldrich #AP164). In the library amplification step, 16 to 21 cycles were used. The purified libraries were quantified by Qubit 1X dsDNA HS Assay Kit and qPCR, analyzed by a Fragment Analyzer System to confirm successful amplification and quality, and sequenced using Novaseq 6000 Sequencing System.

Reads were processed using TrimGalore (v0.6,7) to remove adaptor and low-quality reads with parameters “-q 20, -e 0.1, --length 20, --stringency 1”, and then trimmed reads were mapped to mm10 (GRCm38.p6) reference genome using Bowtie2 (v2.4.2) with parameters “-L 25, -X 2000, --no-mixed --no-discordant”. Unmapped reads and low mapping quality reads (Q<30) were discarded, and PCR duplicates were removed using the function “MarkDuplicates” from Picard tools (v2.27.2). CUT&Tag bigwig tracks were generated using the function “bamcoverage” from Deeptools (v3.5.1) with parameters “-binSize 50 -normSize 2494787188 -exactScaling - extendReads -normalizeUsing RPKM”. CUT&Tag signals on genomic regions were computed using computeMatrix with parameters “-bs 200, --upstream 10000, --downstream 10000”.

### Statistical analysis

No statistical methods were used to predetermine the sample size for experiments. Experiments were not randomized. The investigators were not blinded to allocation during experiments and outcome assessment. Average (mean) and SD were calculated in Excel. Statistical significance is based on unpaired, two-tailed Student’s t test (for absolute values) and two-tailed Fisher’s exact test (for categorical values) calculated in OriginPro or Prism (GraphPad), assuming normal distribution and similar variance. All box plots show median (horizontal black line), mean (small black squares), 25^th^ and 75^th^ percentiles (boxes), 5^th^ and 95^th^ percentiles (whiskers), and 1^st^ and 99^th^ percentiles (crosses). All data were from at least two independent experiments. P values are designated as *P < 0.05, **P < 0.01, ***P < 0.001, and ****P < 0.0001. Nonsignificant values are indicated as N.S.

---

**Figure S1.**
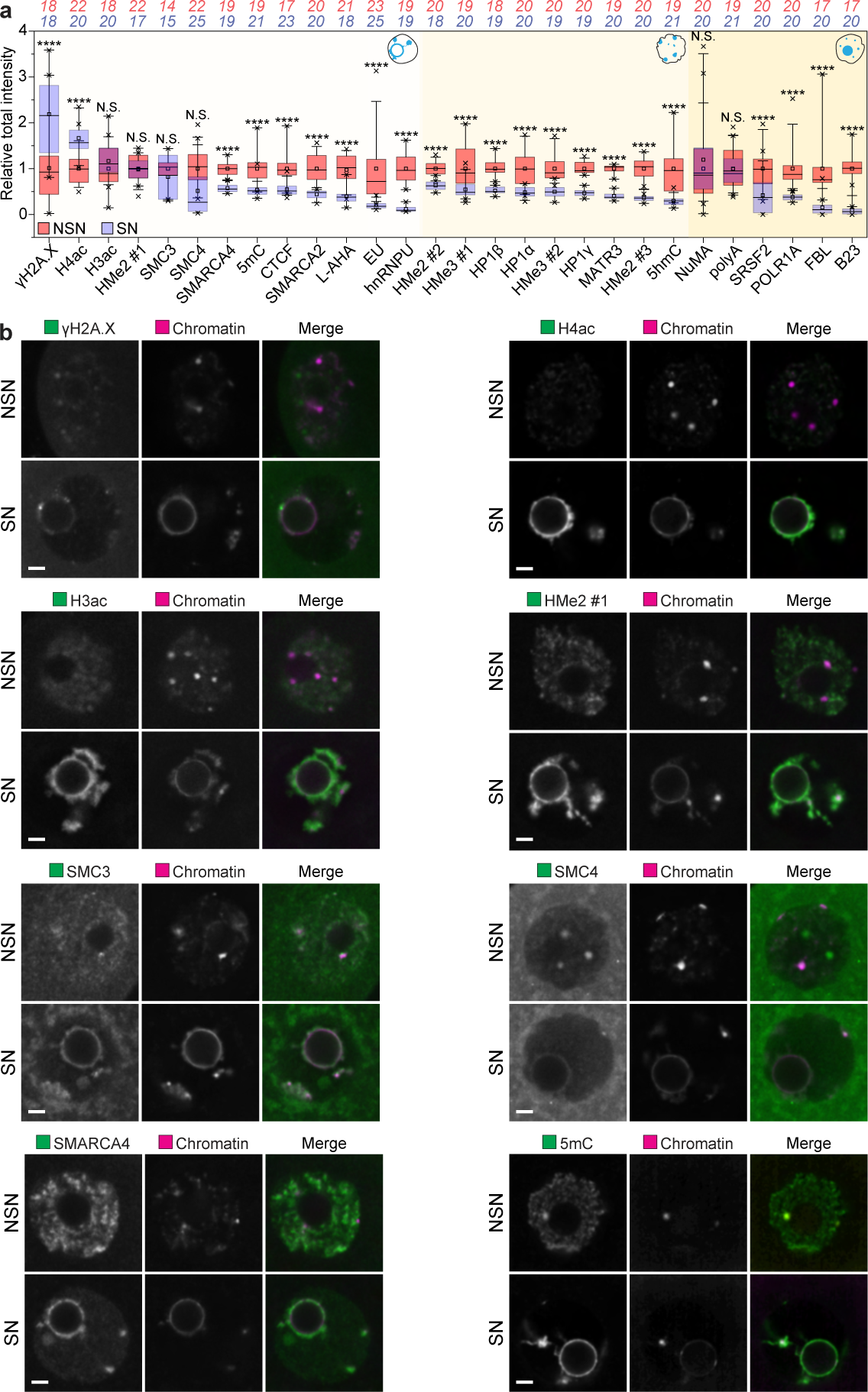
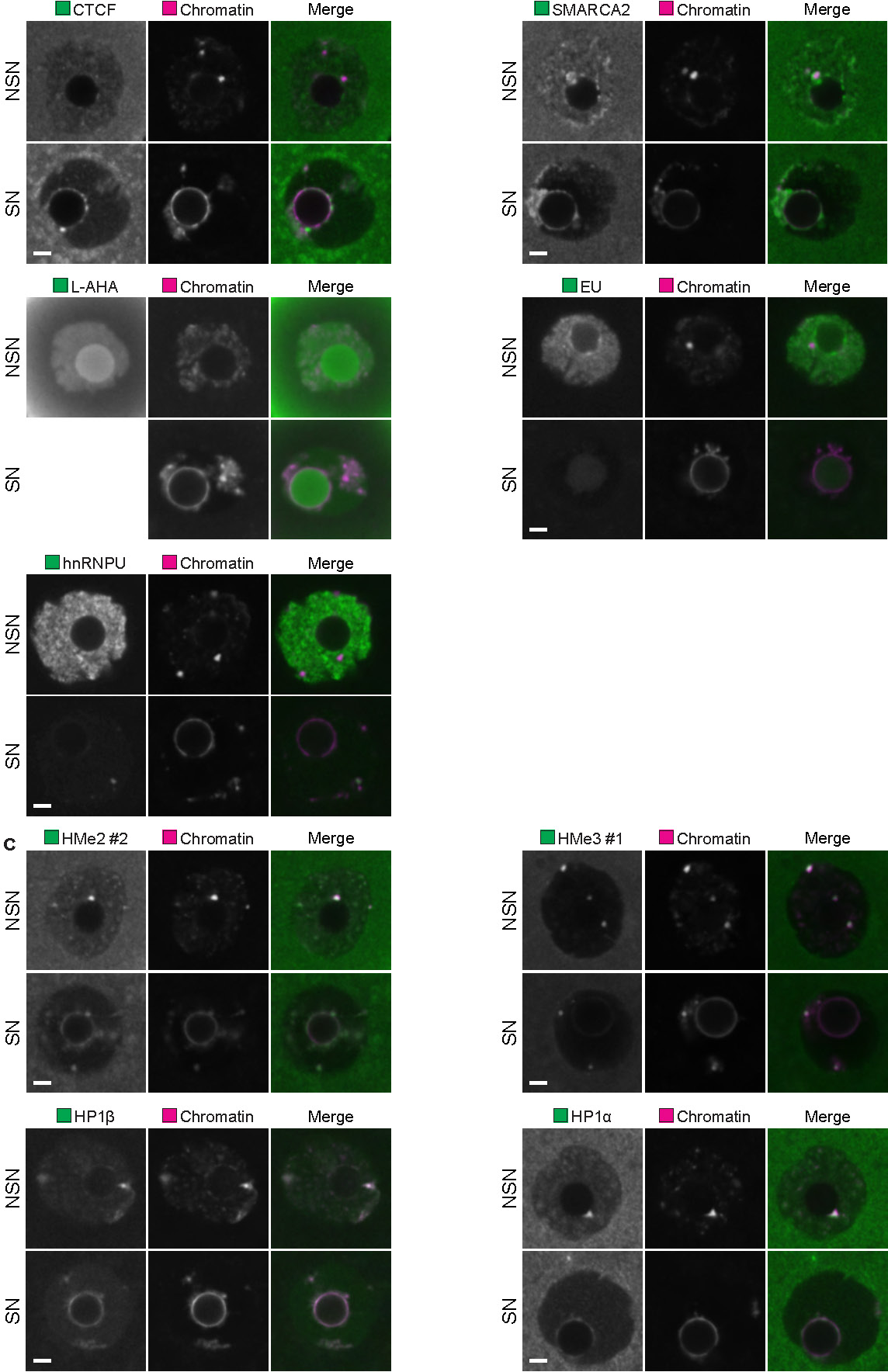
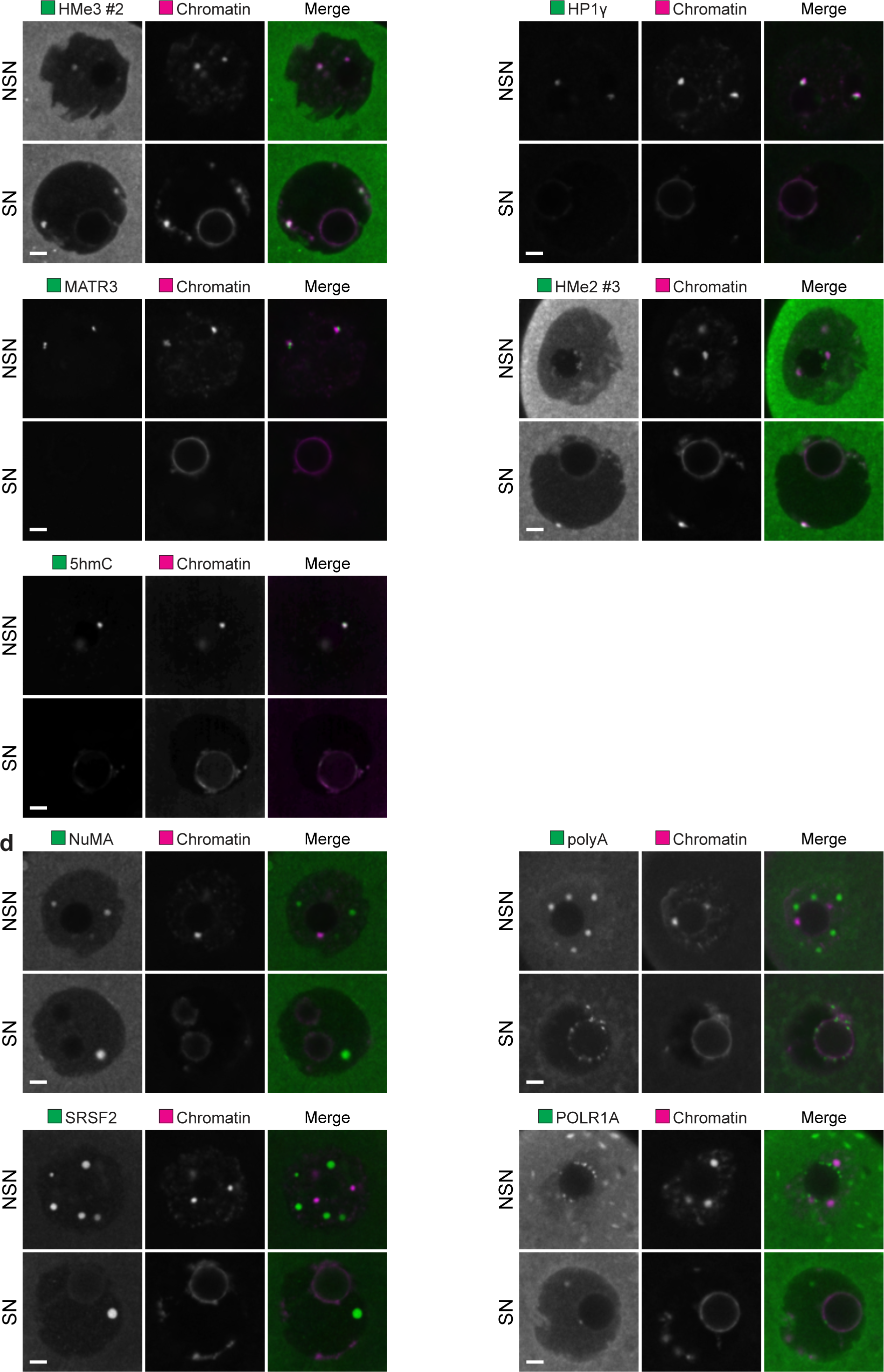
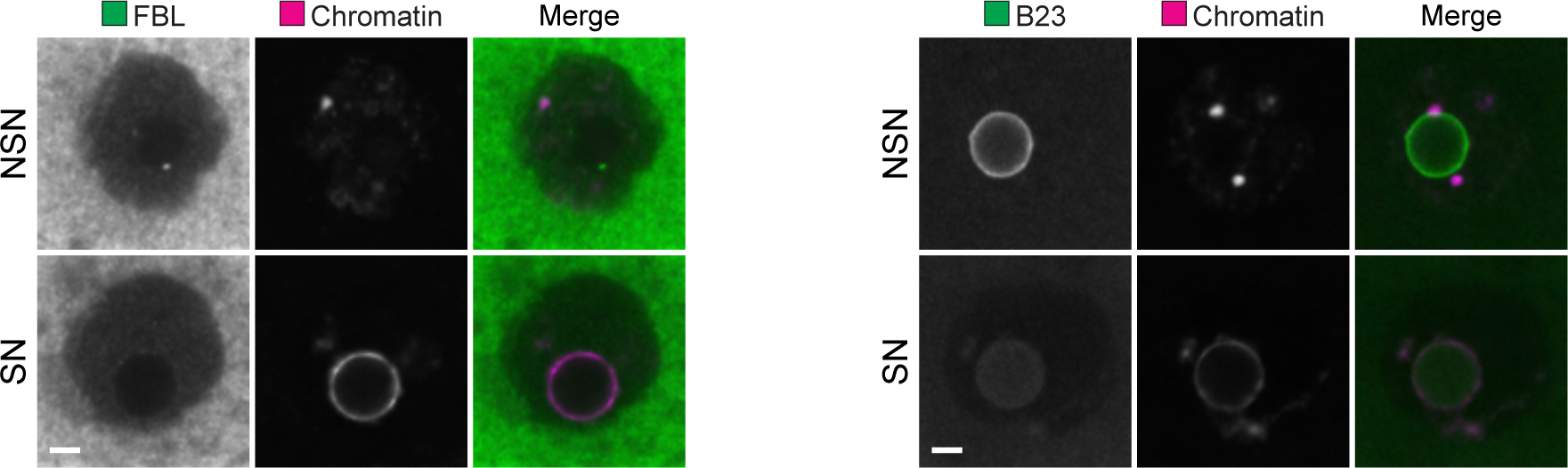
Systematic evaluation of the localization and abundance of common nuclear components in NSN and SN nuclei. **(A)** Automated quantification of total intensity of different nuclear components in NSN and SN oocytes. **(B)** Immunofluorescence images of different nuclear components localizing to chromatin in NSN and SN oocytes. Green, protein of interest; magenta, chromatin (Hoechst). **(C)** Immunofluorescence images of different nuclear components localizing to chromocenters in NSN and SN oocytes. Green, protein of interest; magenta, chromatin (Hoechst). **(D)** Immunofluorescence images of different nuclear components localizing to nuclear bodies in NSN and SN oocytes. Green, protein of interest; magenta, chromatin (Hoechst). ****P < 0.0001. N.S., not significant. See Methods for box plot specifications. The number of oocytes analyzed is specified in italics. Scale bars are 5 µm.

**Figure S2.**
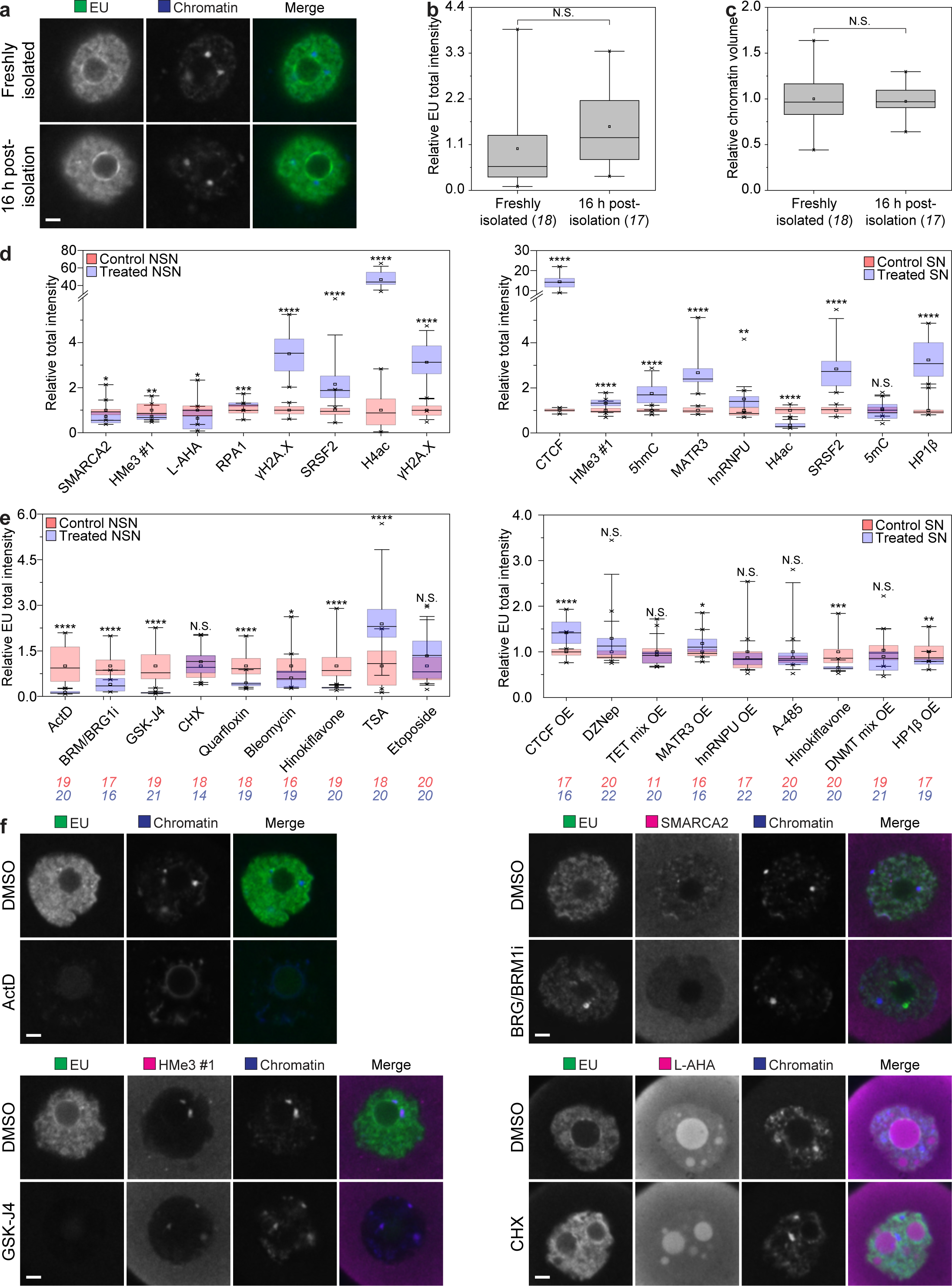
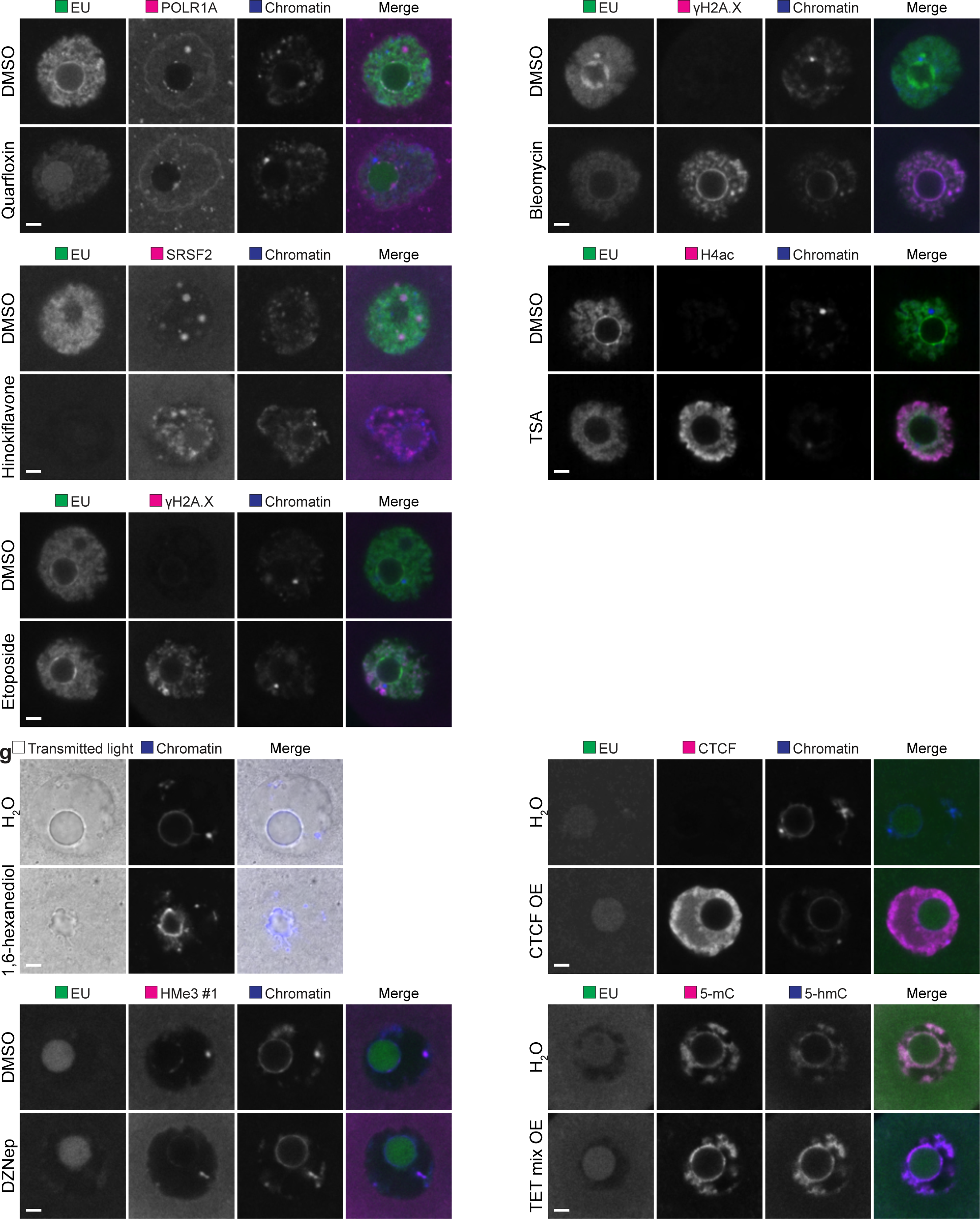
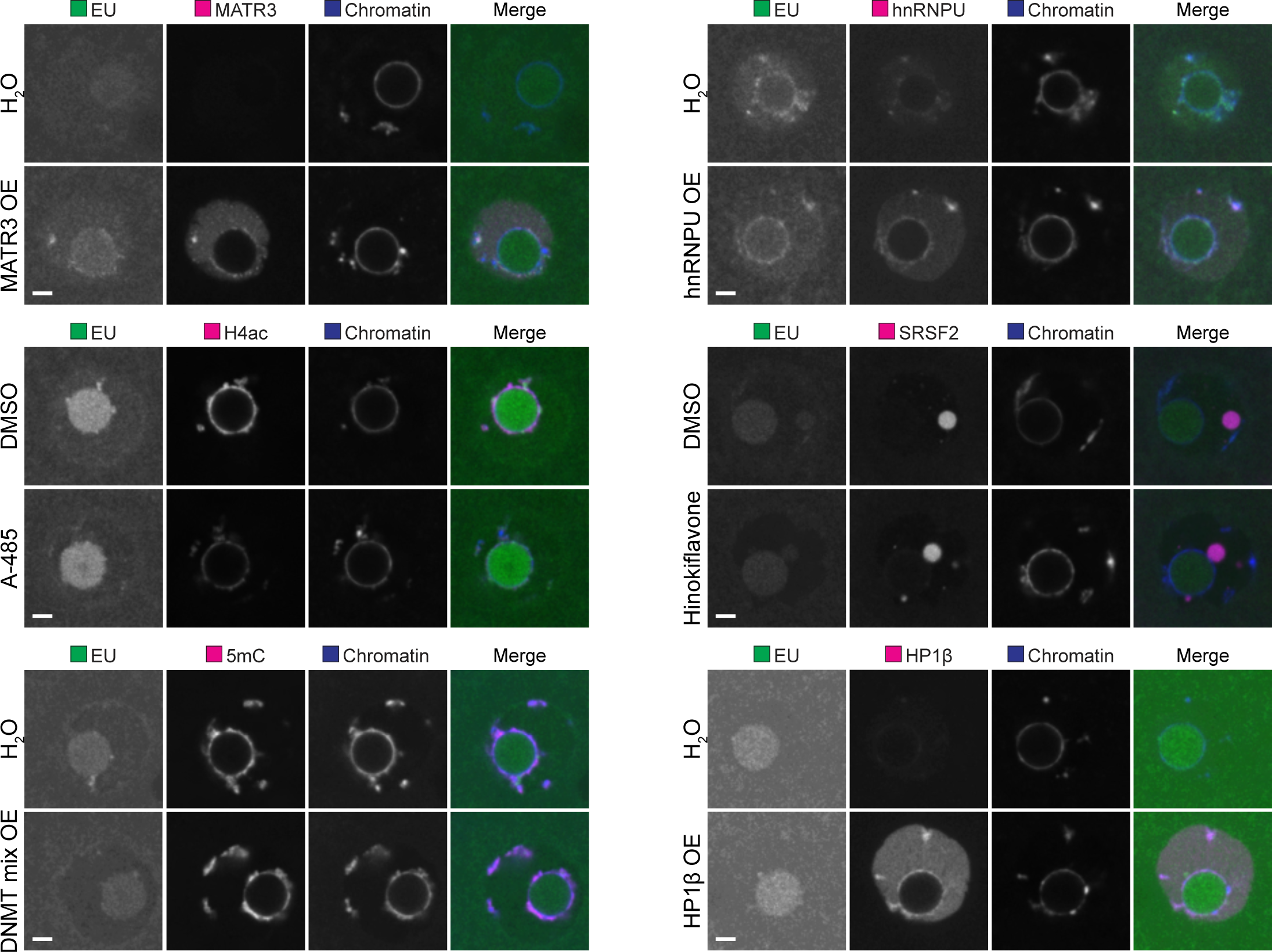
Systematic identification of nuclear components potentially driving NSN-to-SN transition. **(A)** Fluorescence images of NSN oocytes freshly isolated or incubated for 16 h. Green, EU; blue chromatin (Hoechst). **(B and C)** Automated quantification of total intensity of EU and chromatin volume in NSN oocytes freshly isolated or incubated for 16 h. **(D and E)** Automated quantification of total intensity of the target and EU in NSN and SN oocytes treated with different small molecule-inhibitors or overexpressing different nuclear components. **(F)** Immunofluorescence images of NSN oocytes treated with different small-molecule inhibitors. Green, EU; magenta, target; blue, chromatin (Hoechst). **(G)** Immunofluorescence images of SN oocytes treated with different small-molecule inhibitors or overexpressing different nuclear components. Green, EU; magenta or gray, target; blue, chromatin (Hoechst). *P < 0.05. **P < 0.01. ***P < 0.001. ****P < 0.0001. N.S., not significant. See Materials and methods for box plot specifications. The number of oocytes analyzed is specified in italics. Scale bars are 5 µm.

**Figure S3.**
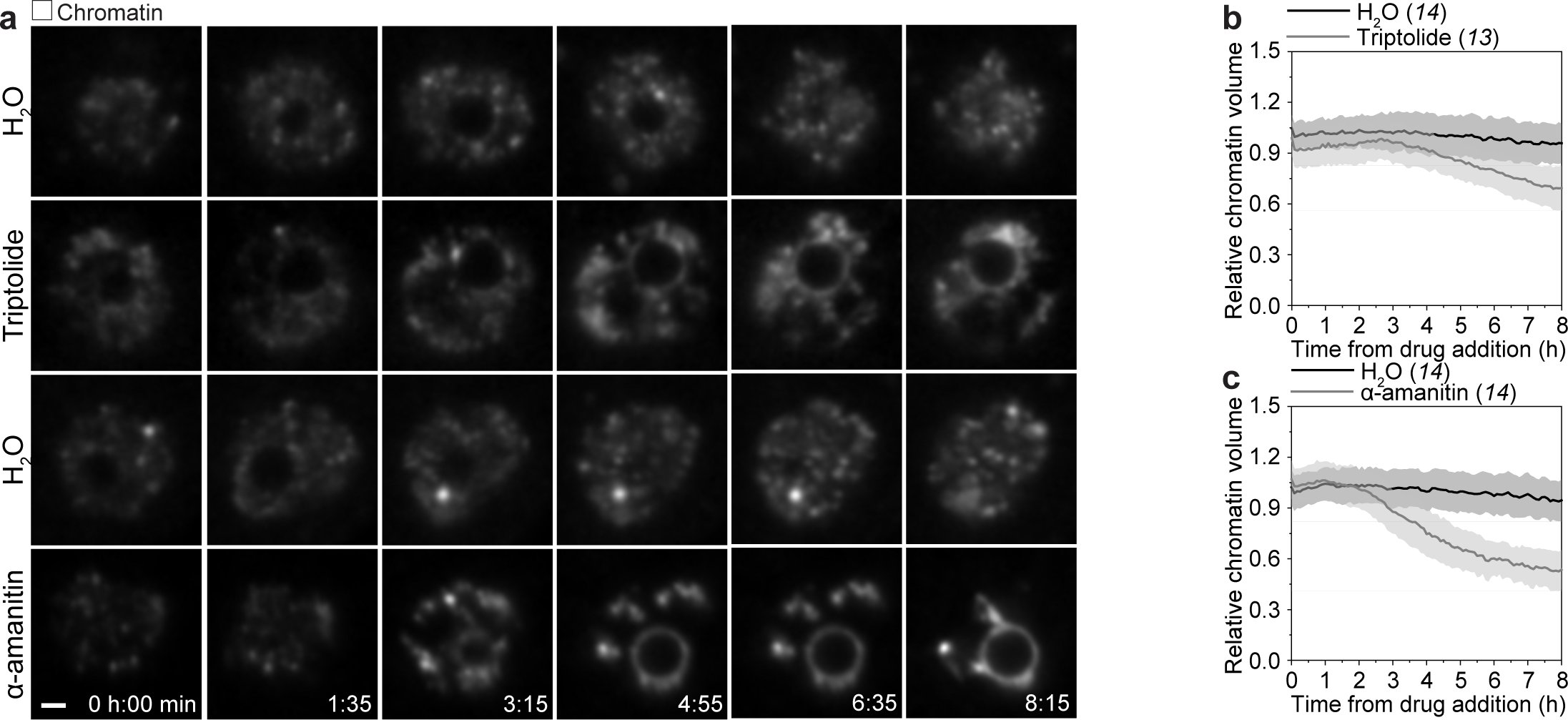
RNAPII inhibitors swiftly induce NSN-to-SN transition in oocytes. **(A)** Still images from time-lapse movies of NSN oocytes treated with triptolide or α-amanitin. Time is given as hours:minutes after drug addition. **(B and C)** Automated quantification of chromatin volume in NSN oocytes treated with triptolide or α-amanitin over time. The number of oocytes analyzed is specified in italics. Scale bars are 5 µm.

**Figure S4.**
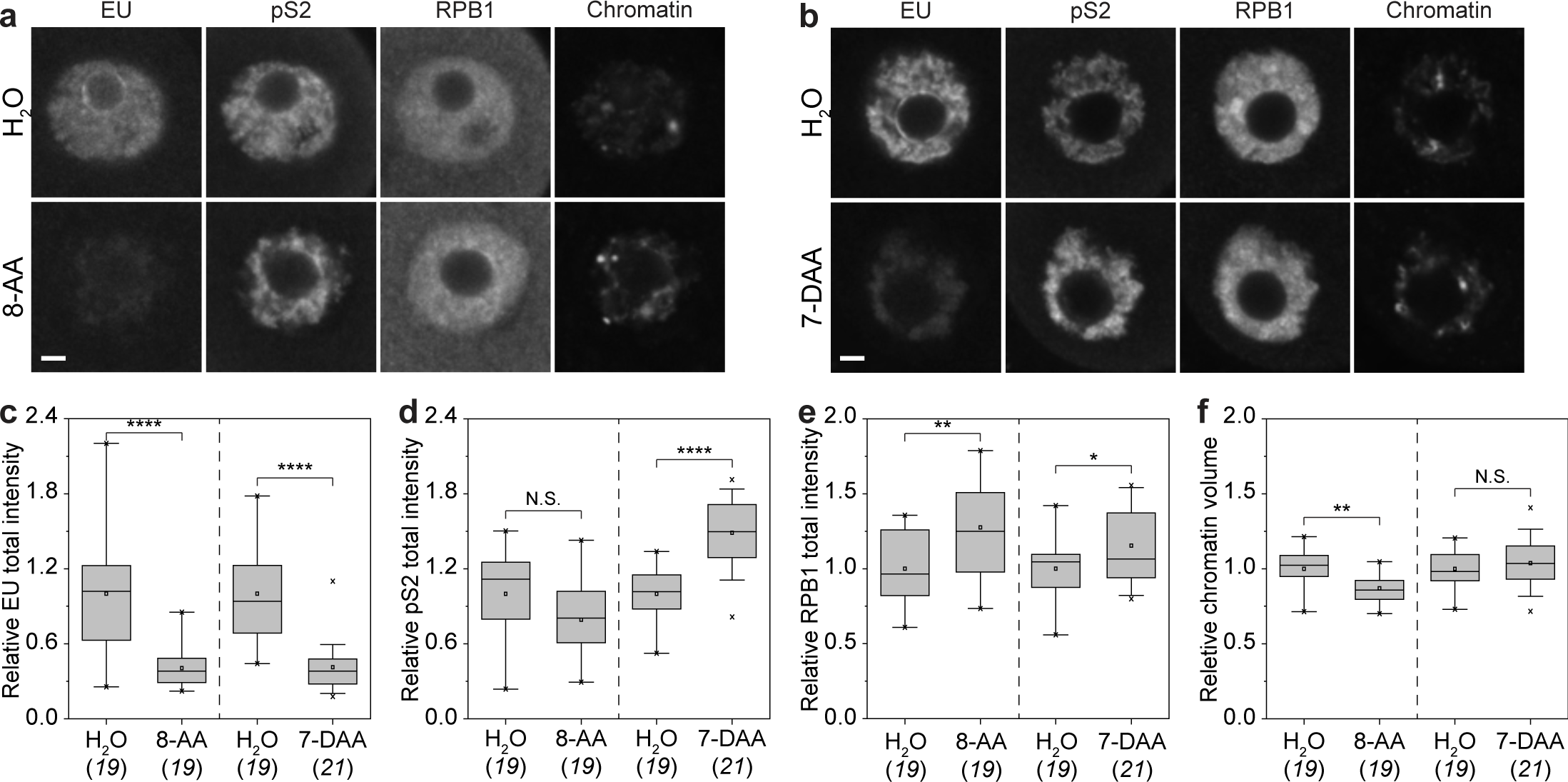
Transcriptional silencing is insufficient to induce NSN-to-SN transition in oocytes. **(A and B)** Immunofluorescence images of NSN oocytes treated with 8-AA or 7-DAA. **(C to F)** Automated quantification of total intensity of EU, pS2, RPB1 and chromatin volume in NSN oocytes treated with 8-AA or 7-DAA. *P < 0.05. **P < 0.01. ****P < 0.0001. N.S., not significant. See Materials and methods for box plot specifications. The number of oocytes analyzed is specified in italics. Scale bars are 5 µm.

**Figure S5.**
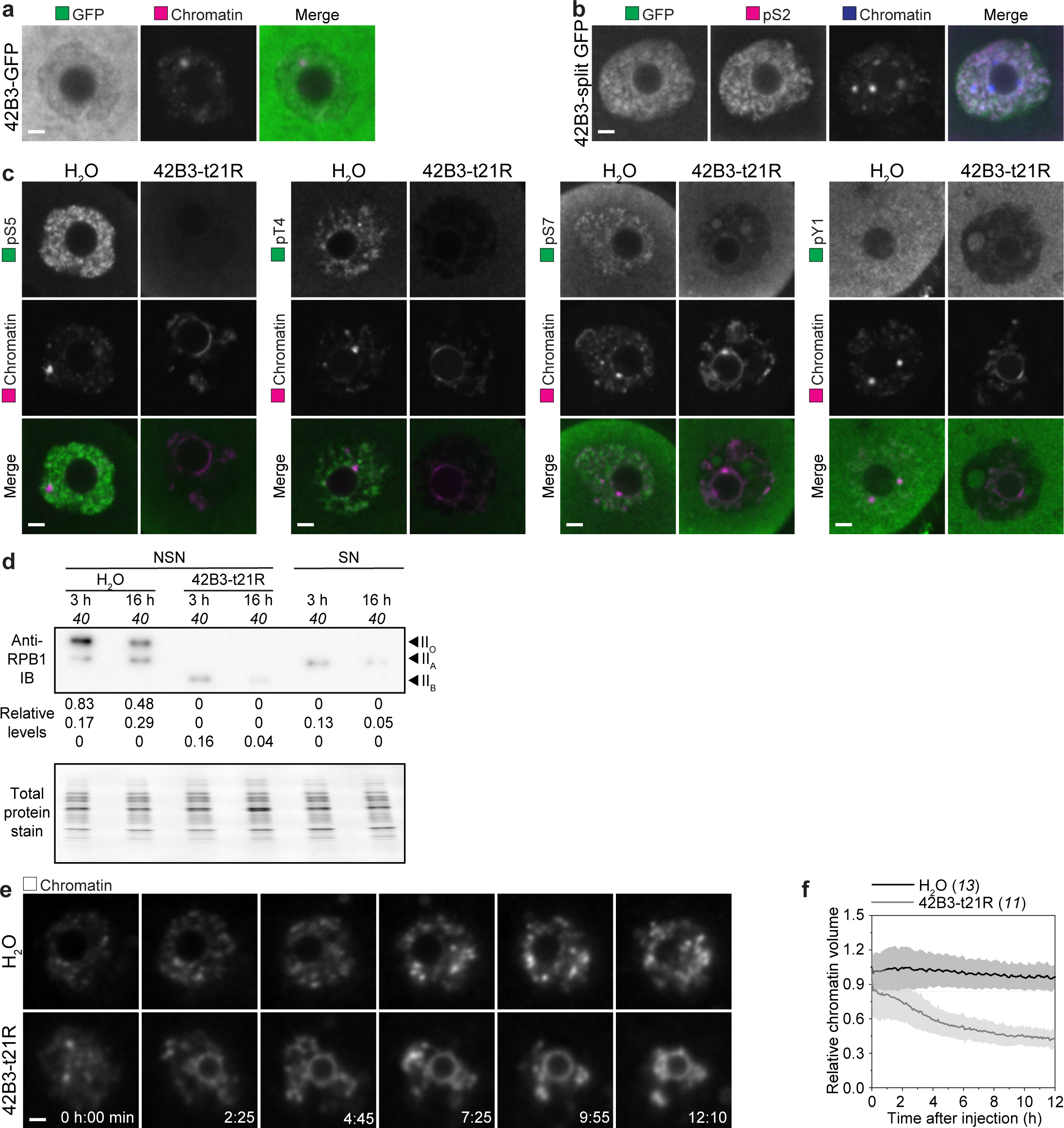
Trim-Away of RNAPII with 42B3-t21R. **(A)** Immunofluorescence images of NSN oocytes expressing 42B3-GFP. Green, GFP (anti-GFP); magenta, chromatin (Hoechst). **(B)** Immunofluorescence images of NSN oocytes expressing 42B3-split GFP. Green, GFP (anti-GFP); magenta, pS2; blue, chromatin (Hoechst). **(C)** Immunofluorescence images of NSN oocytes expressing 42B3-t21R for 3 h. Green, pS5, pT4, pS7 or pY1; magenta, chromatin (Hoechst). **(D)** Immunoblots of control NSN, 42B3-t21R-expressing NSN and SN oocytes cultured for 3 or 16 h. Black arrows mark phosphorylated (IIO), unphosphorylated (IIA) and tailless (IIB) RPB1. The full blot was stained for total protein before immunoblotting. **(E)** Still images from time-lapse movies of NSN oocytes expressing 42B3-t21R. Time is given as hours:minutes from the start of imaging immediately after injection. **(F)** Automated quantification of chromatin volume in NSN oocytes expressing 42B3-t21R over time. The number of oocytes analyzed or loaded is specified in italics. Scale bars are 5 µm.

**Figure S6.**
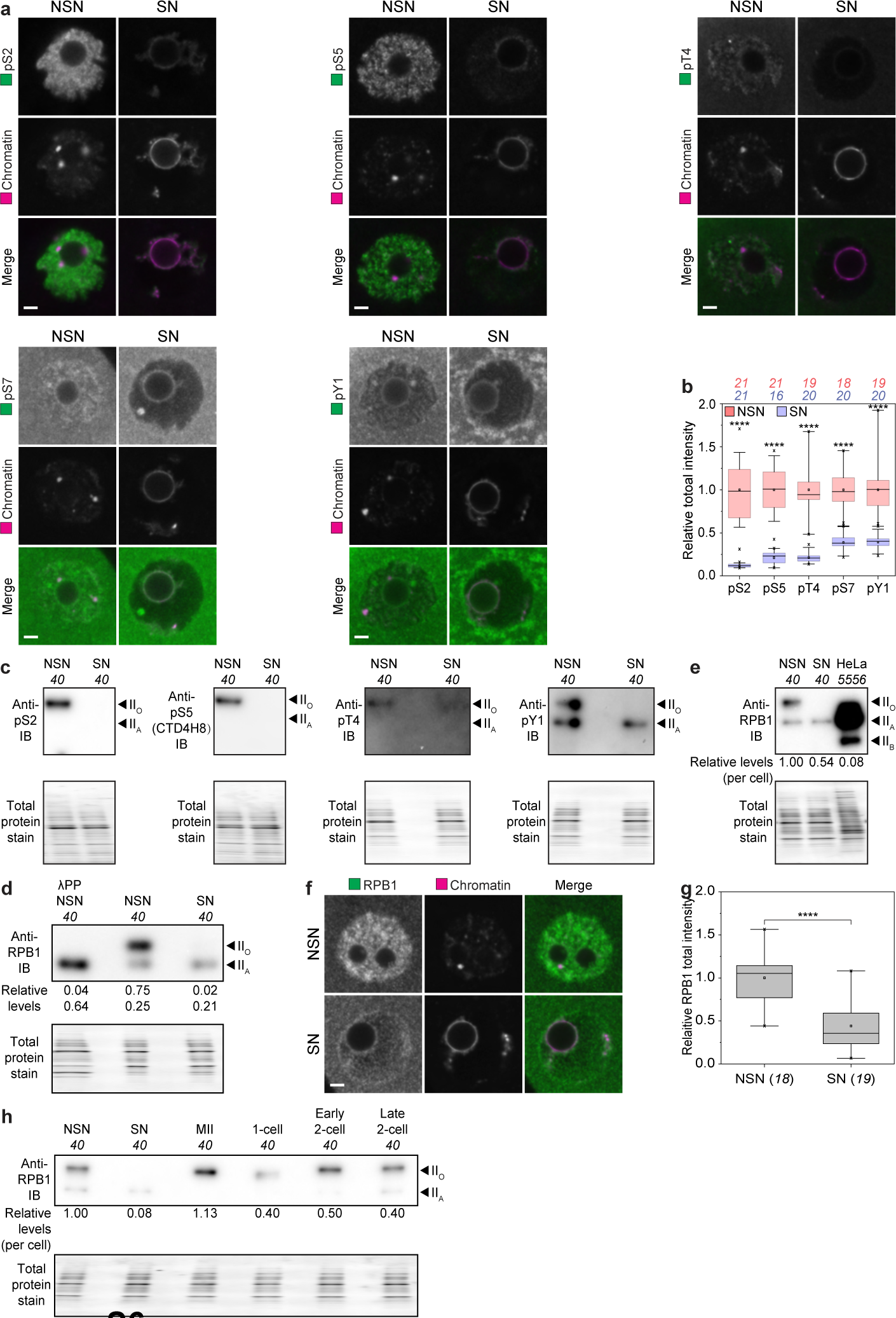
Depletion of phosphorylated RNAPII at protein level during NSN-to-SN transition in vivo. **(A)** Immunofluorescence images of NSN and SN oocytes. Green, pS2, pS5, pT4, pS7 or pY1; magenta, chromatin (Hoechst). **(B)** Automated quantification of total intensity of pS2, pS5, pT4, pS7 and pY1 in NSN and SN oocytes. **(C)** Immunoblots of NSN and SN oocytes. Black arrows mark phosphorylated (IIO) and unphosphorylated (IIA) RPB1. Full blots were stained for total protein before immunoblotting. **(D)** Immunoblots of lambda protein phosphatase (λPP)-treated NSN, NSN and SN oocytes. Black arrows mark phosphorylated (IIO) and unphosphorylated (IIA) RPB1. The full blot was stained for total protein before immunoblotting. **(E)** Immunoblots of NSN, SN oocytes and HeLa cells. Black arrows mark phosphorylated (IIO), unphosphorylated (IIA) and tailless (IIB) RPB1. The full blot was stained for total protein before immunoblotting. **(F)** Immunofluorescence images of NSN and SN oocytes. Green, RPB1; magenta, chromatin (Hoechst). **(G)** Automated quantification of total intensity of RPB1 in NSN and SN oocytes. **(H)** Immunoblots of NSN, SN, MII oocytes, 1-cell, early and late 2-cell embryos. Black arrows mark phosphorylated (IIO) and unphosphorylated (IIA) RPB1. Full blots were stained for total protein before immunoblotting. ****P < 0.0001. The number of oocytes analyzed or loaded is specified in italics. Scale bars are 5 µm.

**Figure S7.**
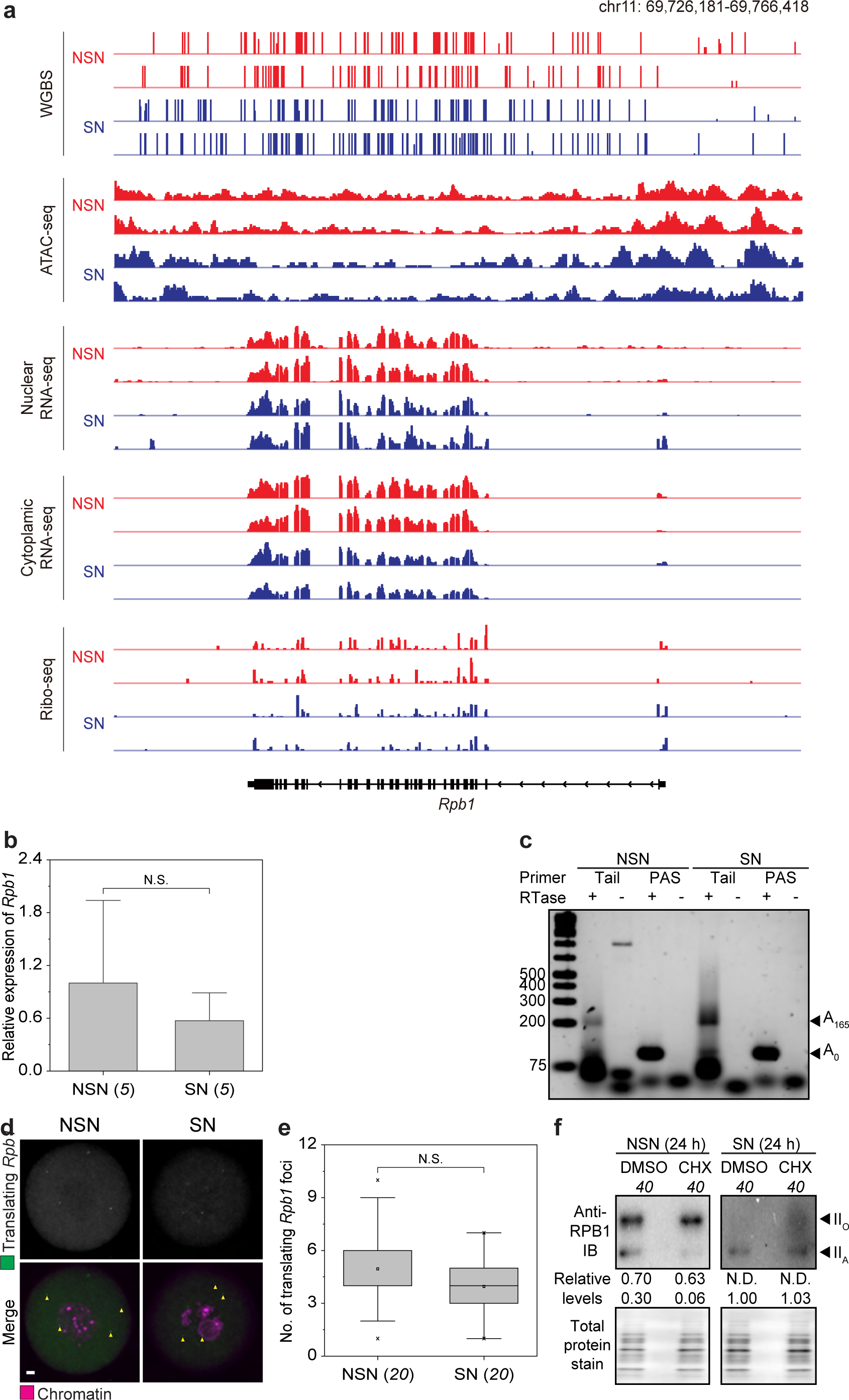
Post-translational depletion of RNAPII during NSN-to-SN transition in vivo. **(A)** IGV browser views showing DNA methylation levels, chromatin accessibility, nuclear RNA-seq (as a proxy for actively transcribing RNAs), cytoplasmic RNA-seq (as a proxy for stored RNAs), Ribo-seq signals for *Rpb1* in NSN and SN oocytes. **(B)** RT-qPCR results showing the expression of *Rpb1* in NSN and SN oocytes. **(C)** Measurement of poly(A) tail length of Rpb1 in NSN and SN oocytes. **(D)** RIBOmap images of translating *Rpb1* in NSN and SN oocytes. Yellow arrowheads highlight translating *Rpb1* foci. **(E)** Automated quantification of the number of translating *Rpb1* foci in NSN and SN oocytes. **(F)** Immunoblots of NSN and SN oocytes treated with CHX for 24 h. Black arrows mark phosphorylated (IIO) and unphosphorylated (IIA) RPB1. Full blots were stained for total protein before immunoblotting. N.D., not determined. N.S., not significant. See Materials and methods for box plot specifications. The number of oocytes analyzed or loaded is specified in italics. Scale bars are 5 µm.

**Figure S8.**
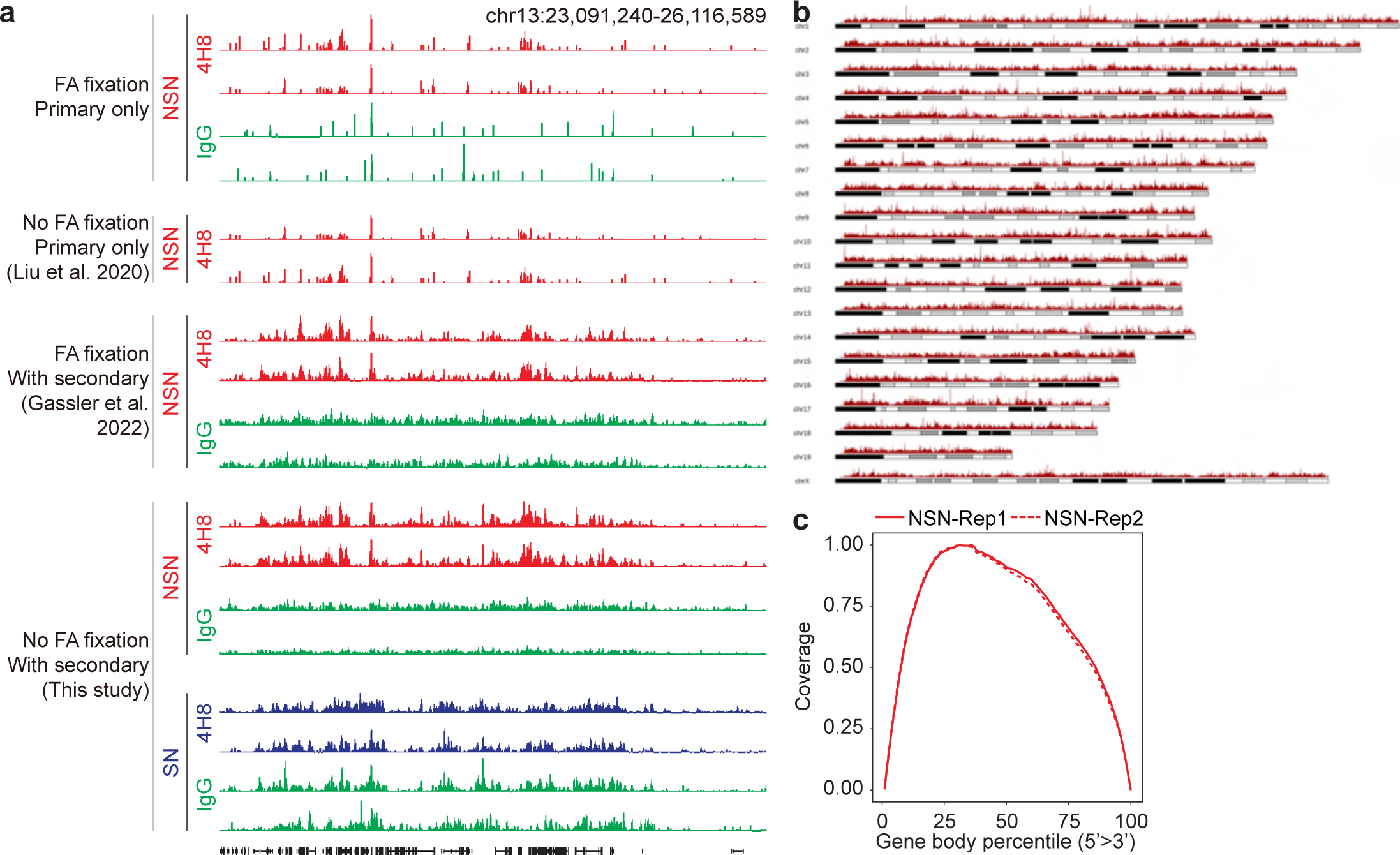
Optimizations of CUT&Tag for low inputs and additional bioinformatics analyses. **(A)** IGV browser views showing CUT&Tag libraries generated by different protocols with anti-CTD pS5 (4H8). No IgG libraries could be obtained with the protocol with no formaldehyde (FA) fixation and with primary antibody only. Only the protocol with no FA fixation and with secondary antibody amplification yielded IgG and 4H8 libraries for both NSN and SN oocytes. **(B)** Ideogram showing pS2 CUT&Tag signals on different chromosomes in NSN oocytes. **(C)** Gene body coverage to show no 3’ bias in the NSN nuclear RNA-seq data.

**Figure S9.**
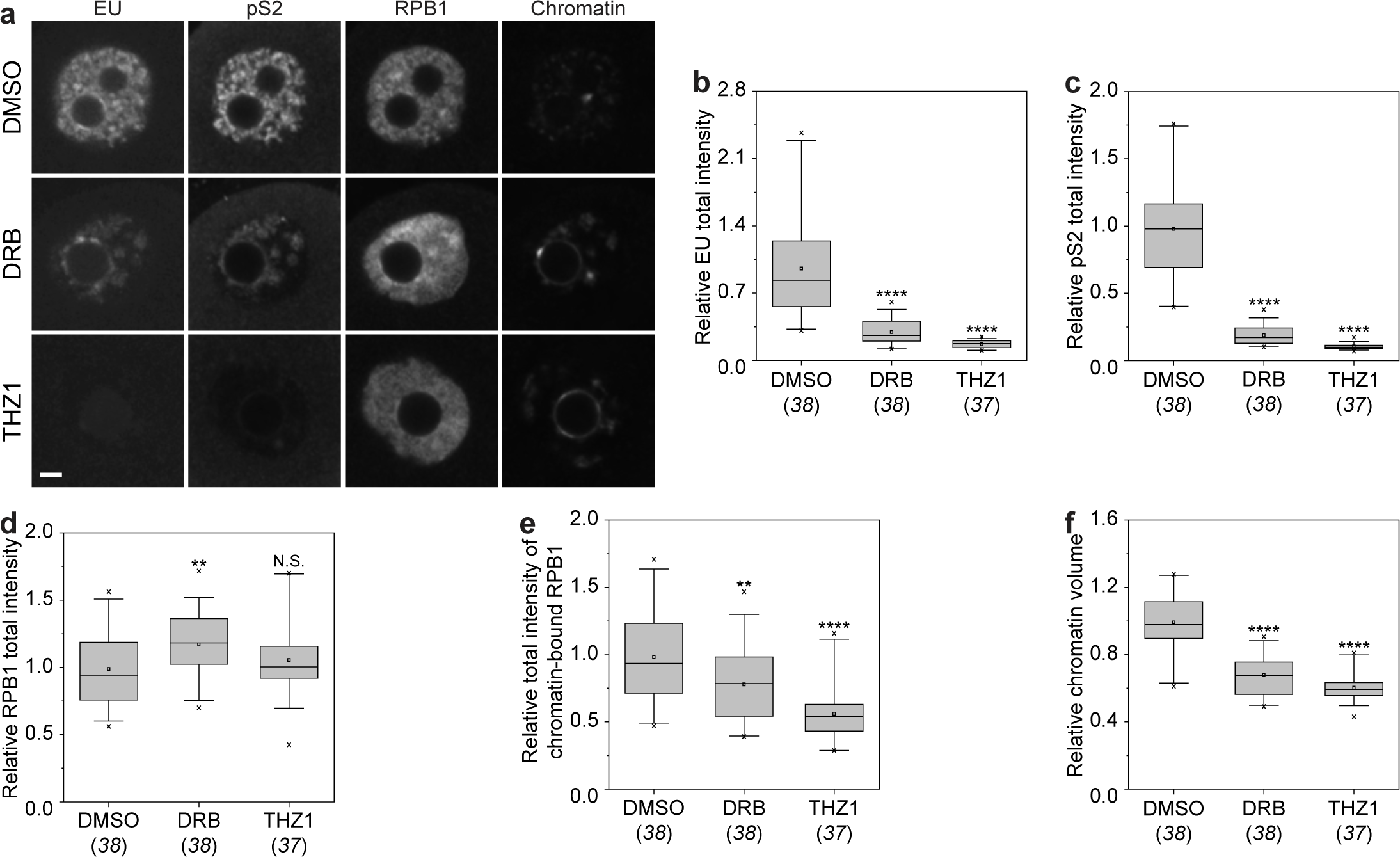
Releasing chromatin-bound RNAPII is sufficient to induce NSN-to-SN transition in oocytes. **(A)** Immunofluorescence images of NSN oocytes treated with DRB or THZ1. **(B to F)** Automated quantification of total intensity of EU, pS2, RPB1, chromatin-bound RPB1 and chromatin volume in NSN oocytes treated with DRB or THZ1. *P < 0.05. **P < 0.01. ****P < 0.001. N.S., not significant. See Materials and methods for box plot specifications. The number of oocytes analyzed is specified in italics. Scale bars are 5 µm.

**Table S1.**
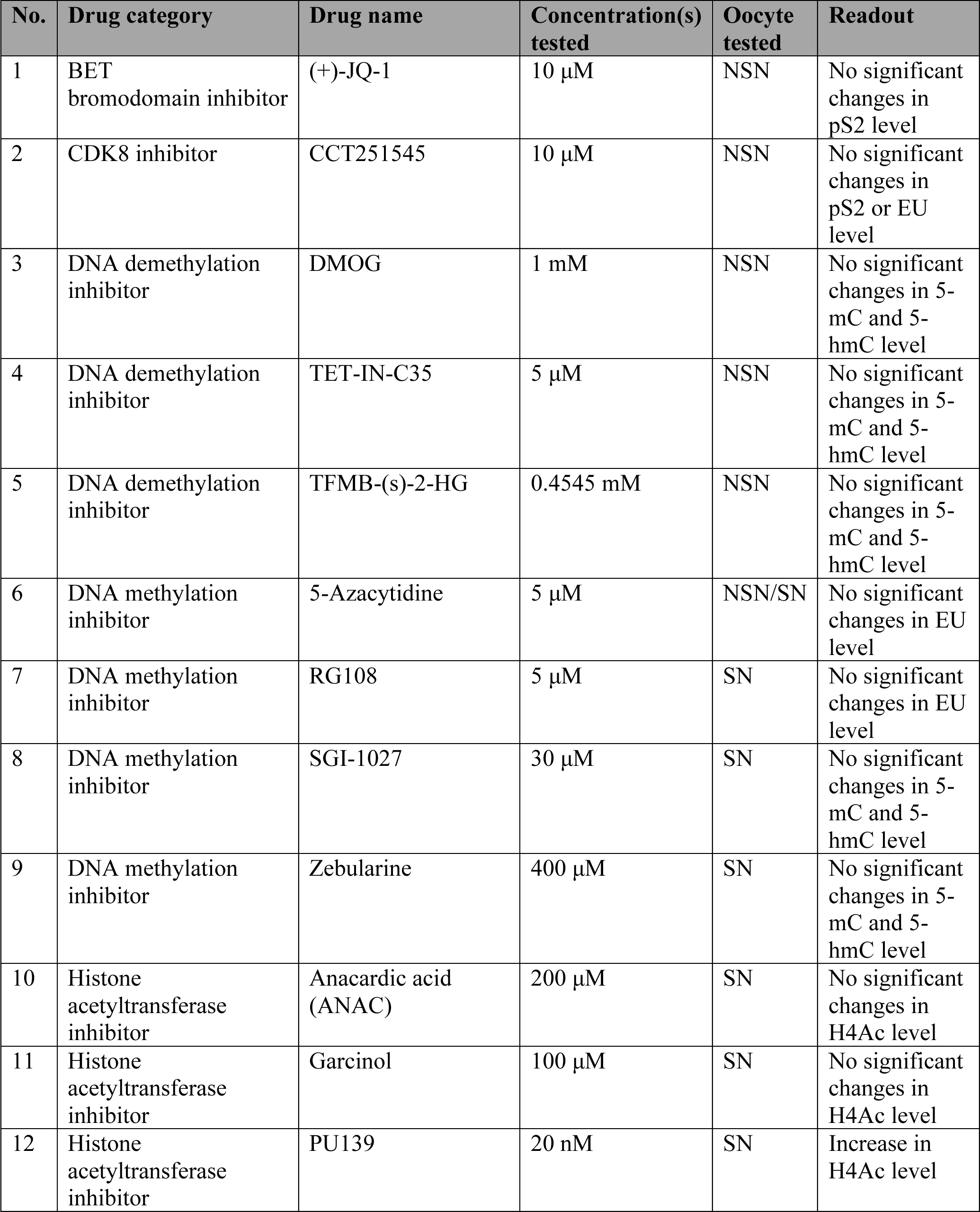
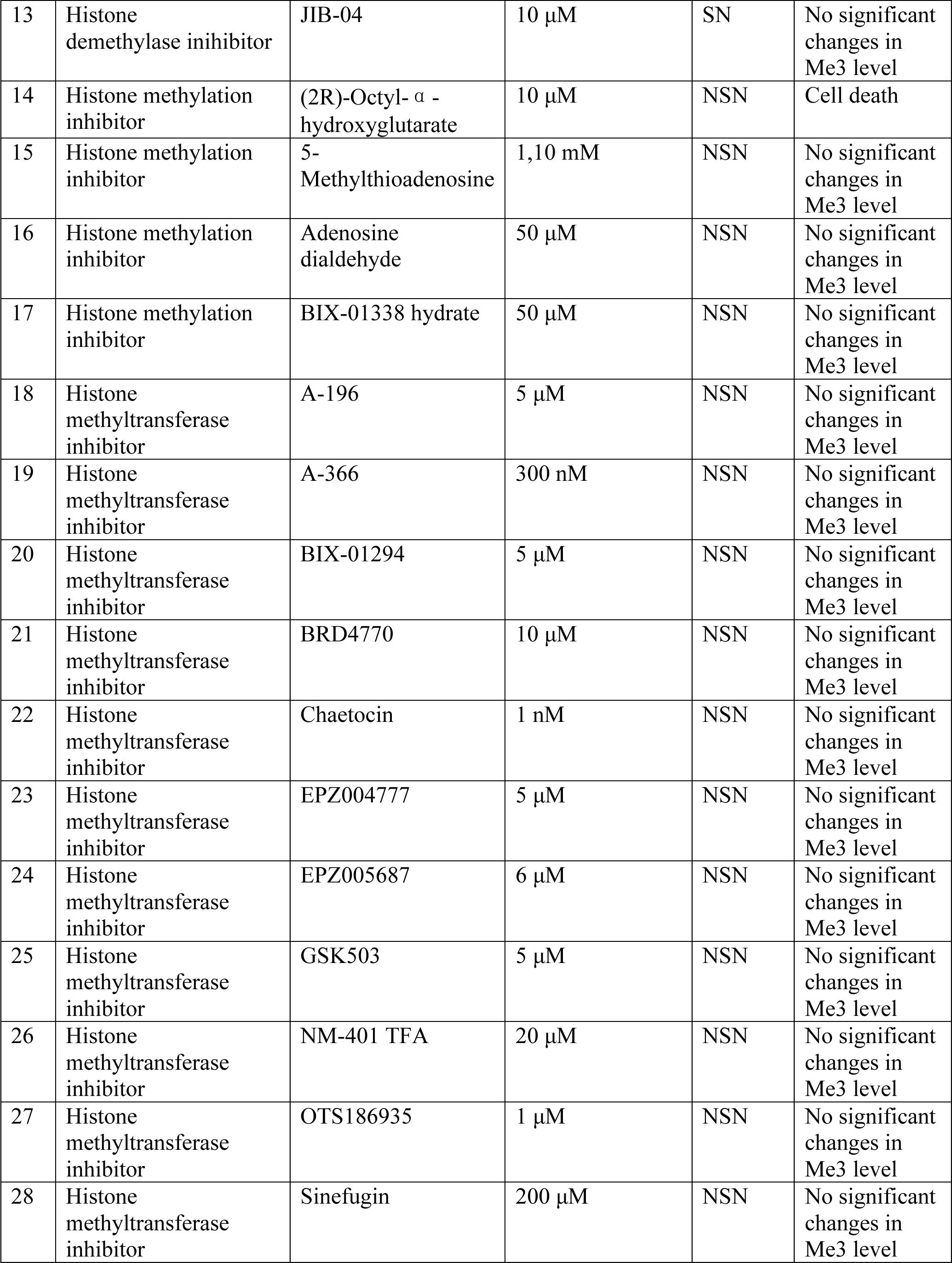
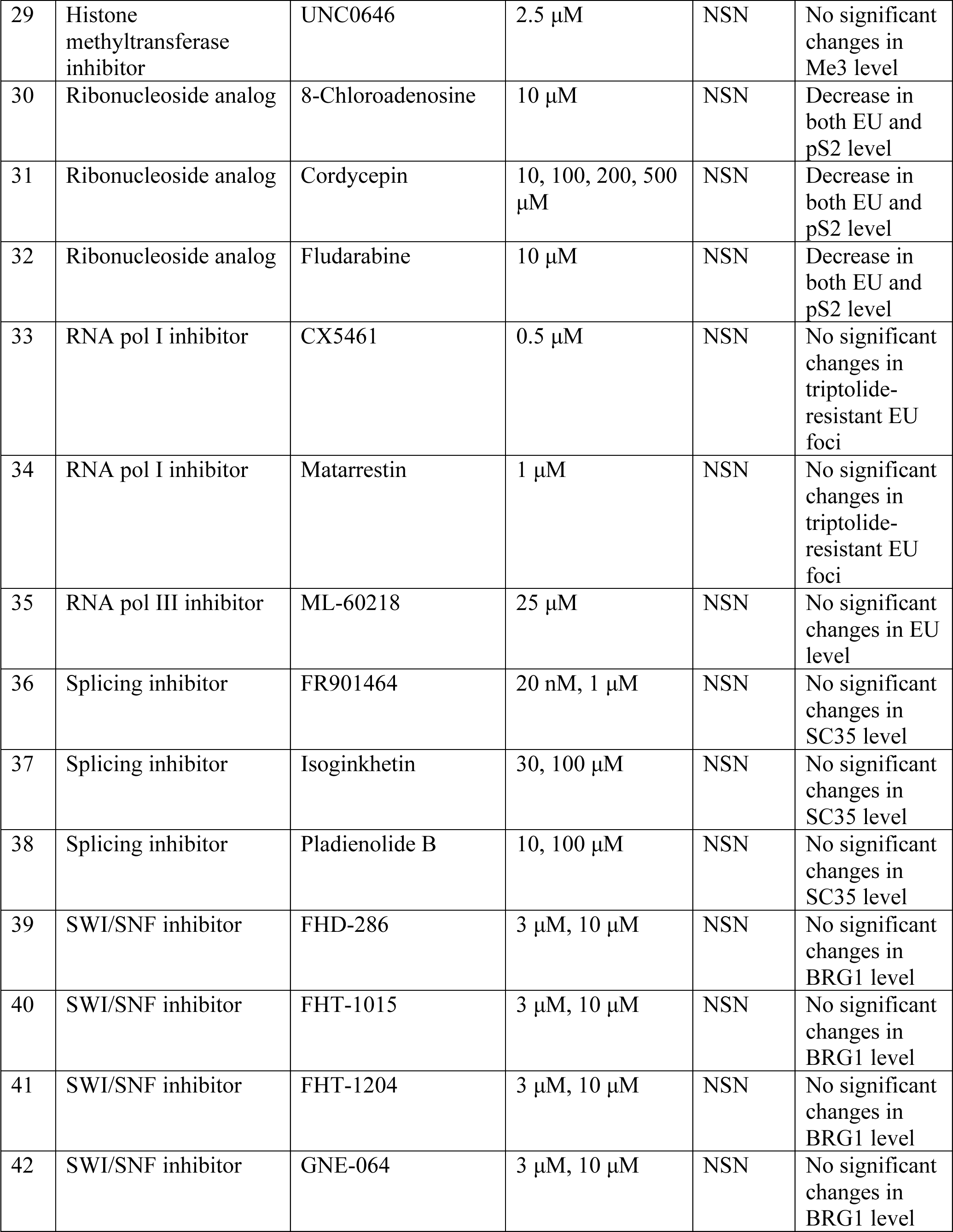
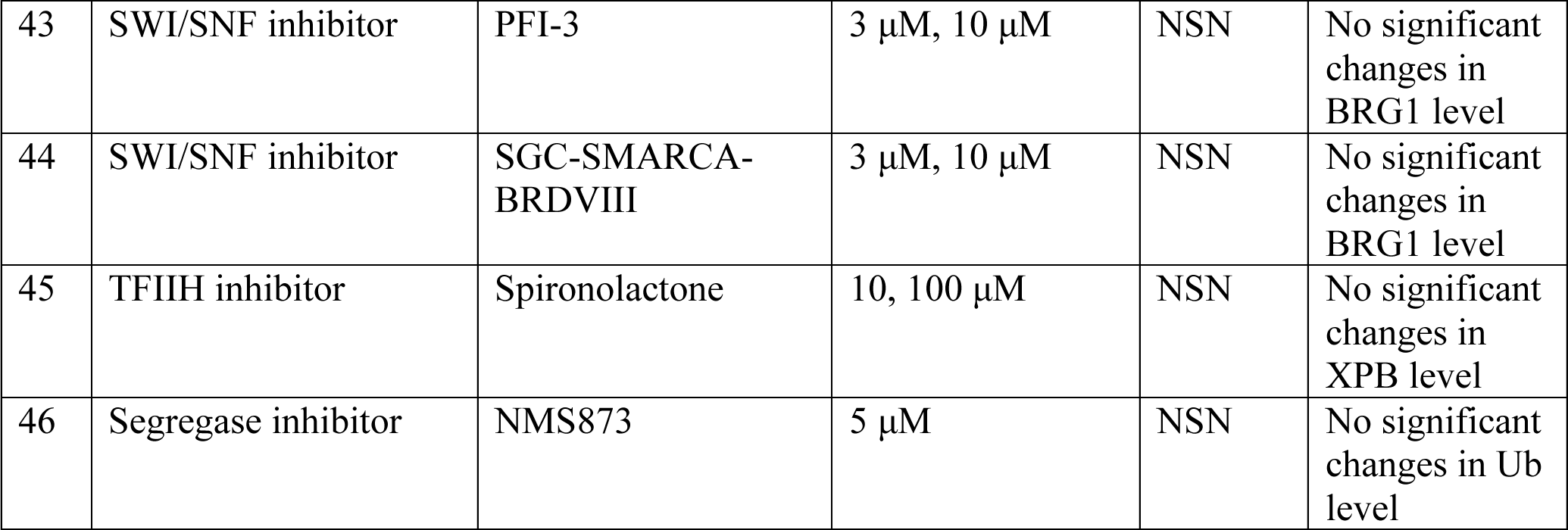
List of tested small-molecule inhibitors that did not work in this study.

**Video S1**

Time-lapse movie of a water-treated NSN oocyte stained with SPY555-DNA (gray, to label chromatin).

**Video S2**

Time-lapse movie of a triptolide-treated NSN oocyte stained with SPY555-DNA (gray, to label chromatin).

**Video S3**

Time-lapse movie of a water-treated NSN oocyte stained with SPY555-DNA (gray, to label chromatin).

**Video S4**

Time-lapse movie of an α-amanitin-treated NSN oocyte stained with SPY555-DNA (gray, to label chromatin).

**Video S5**

Time-lapse movie of a control NSN oocyte stained with SPY555-DNA (gray, to label chromatin).

**Video S6** Time-lapse movie of a NSN oocyte expressing 42B3-t21R and stained with SPY555-DNA (gray, to label chromatin).

**Video S7** Time-lapse movie of a control growing human oocyte stained with SPY555-DNA (gray, to label chromatin).

**Video S8** Time-lapse movie of a growing human oocyte expressing 42B3-t21R and stained with SPY555-DNA (gray, to label chromatin).

